# Enemies, more than sex, shape butterfly post-mating odor evolution

**DOI:** 10.64898/2026.05.31.729122

**Authors:** Liana O. Greenberg, Qi Wang, Jonne Bonnet, Esther L. te Lindert-Blommert, Jip van Buul, Tjomme Fernhout, Julia Friman, Niek Palmen, Alex Rozing, Esther van Tatenhove, Jonne Veldboom, Astrid T. Groot, Kees van Oers, M. Eric Schranz, Bas J. Zwaan, Alexander Haverkamp, Bart A. Pannebakker, Nina E. Fatouros

## Abstract

Not all odors influencing mating behavior evolve as sex pheromones. Female butterflies’ post-mating odors have been considered species-specific anti-aphrodisiac pheromones shaped by sexual selection but may also serve broader ecological roles shaped by natural selection. Males transfer odors to females that repel rivals, yet the widespread use of these compounds across phyla makes them targets for eavesdropping, such as by phoretic egg parasitoids. We show that in cabbage white butterflies (*Pieris* spp.), these odors are highly variable and attract parasitoids, deter predators, and influence oviposition. Using gas chromatography and electroantennography, we demonstrate that odor emission and perception lack species-specificity: compounds once thought unique to *P. brassicae* and *P. rapae* are shared across Pieridae. In *P. napi*, odor variation among populations correlates with parasitoid pressure, but not with latitude, genetic distance, or mating frequency, suggesting ecological rather than sexual drivers. In *P. brassicae*, CRISPR/Cas9 disruption of odor perception alters oviposition and increases susceptibility to parasitism. Moreover, these odors render females unpalatable to birds. Together, our results show that post-mating odors in *Pieris* butterflies may act as aposematic signals. We provide evidence that these signals evolve under multiple selective pressures, balancing deterrence of mates and predators, parasitoid avoidance, and host-plant interactions. These findings suggest that chemical signals should be viewed as integrating ecological and reproductive pressures, rather than being interpreted solely through the lens of sexual communication.

## Main text

Chemosensory cues are essential for species recognition and mate location across animal phyla^1–3^, simultaneously conveying valuable information to potential mates and enemies alike^4,5^. Darwin developed the theory of sexual selection to explain how conspicuous traits, such as the peacock’s tail, can evolve despite imposing survival costs^6^. Sex pheromones epitomize this tension: while they facilitate mate attraction and reproductive isolation^5^, they can also be exploited by predators and parasitoids^7–9^. This dual role has made sex pheromones a central focus in studies of speciation and the balance between sexual and natural selection^6,10^.

Among sex pheromones, post-mating odors are a distinctive class, transferred from males to females during copulation. First described as “post-mating female odors” in *Heliconius erato* (Nymphalidae)^11^, these cues have since been identified in multiple *Pieris* (Pieridae) species and are now commonly termed anti-aphrodisiac pheromones, a type of post-mating sex pheromone that deters additional courtship^12,13^. The three compounds of *Pieris* anti-aphrodisiacs are benzyl cyanide (*P. brassicae*), methyl salicylate (in *P. rapae* and *P. napi*), and indole (in *P. rapae*). Males synthesize these compounds *de novo* from nutrients acquired during feeding^14^. These compounds are transferred to females during copulation via the spermatophore, along with sperm and nutrients^15^. The same compounds occur as products emitted by the Brassicaceae host-plants of *Pieris* butterflies both constitutively^16^ and/or as herbivore^17^ or oviposition-induced volatiles^17,18^.

While these odors signal mating status and repel further courtship, they also expose mated individuals to heightened risk from natural enemies that exploit these cues to locate hosts or prey^19^ or assess prey palatability^20^. These odors act as kairomones, directly attracting parasitoids to butterflies and facilitating phoretic transport to oviposition sites^19,21–23^. They can also induce plant defenses that further recruit parasitoids^17,24–28^.

The rapid diversification of these odors has primarily been attributed to (intra)sexual selection, driven by male competition to secure paternity through increasingly effective repellents^13,29^. Females would then benefit as well through reduced male harassment during oviposition, but this remains unsubstantiated. In *Pieris*, females gain nutrients from remating^30^ and tend not to accept copulation until they are receptive, meaning that males quickly abandon unreceptive females even in the absence of anti-aphrodisiac odors^31^. In addition, in *P. napi*, population-level variation in polyandry does not correspond to differences in spermatophore-derived compounds^32,33^. Rapid population-level divergence has also been observed in the post-mating odors of *Heliconius*^13^ and divergence between species does not correspond with mating system^34^. Thus, the assumed sexual conflict over anti-aphrodisiac use as a driver of evolution of these odors shows limited explanatory power^34–36^. These findings suggest that antagonistic coevolution between sexes alone is unlikely to account for odor variation^37^. Accordingly, we refer to these compounds more broadly as post-mating odors, recognizing their diverse ecological roles and evolutionary pressures.

Signal exploitation by natural enemies may accelerate divergence: when populations face different predators or parasitoids, selection to evade exploitation can drive shifts in signaling traits and reinforce reproductive isolation^7^. Such interactions between natural and sexual selection can promote speciation when mating preferences align with ecologically selected traits^6^. The possibility for natural selection on post-mating odors is especially likely when cues are broadly detectable, attract natural enemies, and confer little direct sexual benefit. In such cases, natural selection may outweigh sexual selection, leading to the reduction or loss of such signals^7,38,39^. Post-mating odors in *Pieris* butterflies provide a compelling example of this dynamic^23,37^.

Despite some evidence of odor species-specificity^12,15^, the components of post-mating odors in *Pieris* butterflies are environmentally widespread, being emitted by both insects and their host-plants^40^. This ubiquity suggests overlap among species and vulnerability to exploitation^39^, with population-level variation in odor profiles possibly shaped more by ecological than reproductive factors^5^. We hypothesized that natural enemies, such as egg parasitoids (*Trichogramma* spp.), exert stronger selection on post-mating odor blends than mates, as these cues can increase the risk of egg parasitism^19,22,23^, and changes in these blends within a species can drive differential responses by *Trichogramma*^41^. Such antagonistic pressures may favor the reduction or modification of odor emission to evade detection^4,7^, while in other contexts the emission of specific compounds, such as benzyl cyanide, may function as aposematic signals that deter predators by signaling chemical defense^20^. Given that these same odors also mediate oviposition behavior and elicit defense responses in host-plants^16–18^, we further hypothesized that disruption of post-mating odor cues could cascade into altered plant–insect interactions. Collectively, these hypotheses predict that post-mating odors mediate diverse ecological interactions and that their evolution is primarily shaped by natural selection, extending well beyond their traditional role in sexual communication.

### Post-mating odor compounds are shared between species

Using direct-contact PDMS fiber sampling^42^, we detected four principal post-mating compounds in mated *P. napi* females, with negligible emission in unmated controls (Fig. 1A, B). Methyl salicylate is the established anti-aphrodisiac in *P. napi*, while benzyl cyanide and indole serve this function in the related species *P. brassicae* and *P. rapae*, respectively^12^. While methyl salicylate alone has been shown to deter male *P. napi*, the behavioral effect of benzyl cyanide and indole in this species remains untested despite their established anti-aphrodisiac role in congeners. We included guaiacol in our chemical analysis because of its prior detection in *P. napi* headspace collections^14^. We did so even though previous studies considered the amount of the compound to be insignificant and likely a by-product of pheromone biosynthesis^14^ because the presence of minor compounds may still have significant behavioral effects^40^, and guaiacol may modulate or amplify the activity of methyl salicylate^43^. Thus, the post-mating odor profile of *P. napi* incorporates compounds known to function as anti-aphrodisiacs in congeners. Benzyl cyanide, entirely absent in virgins, and indole, present only at low levels in virgins, likely share a male-transferred origin, though this has only been established in congeners^12,14^.

**Figure 1.**
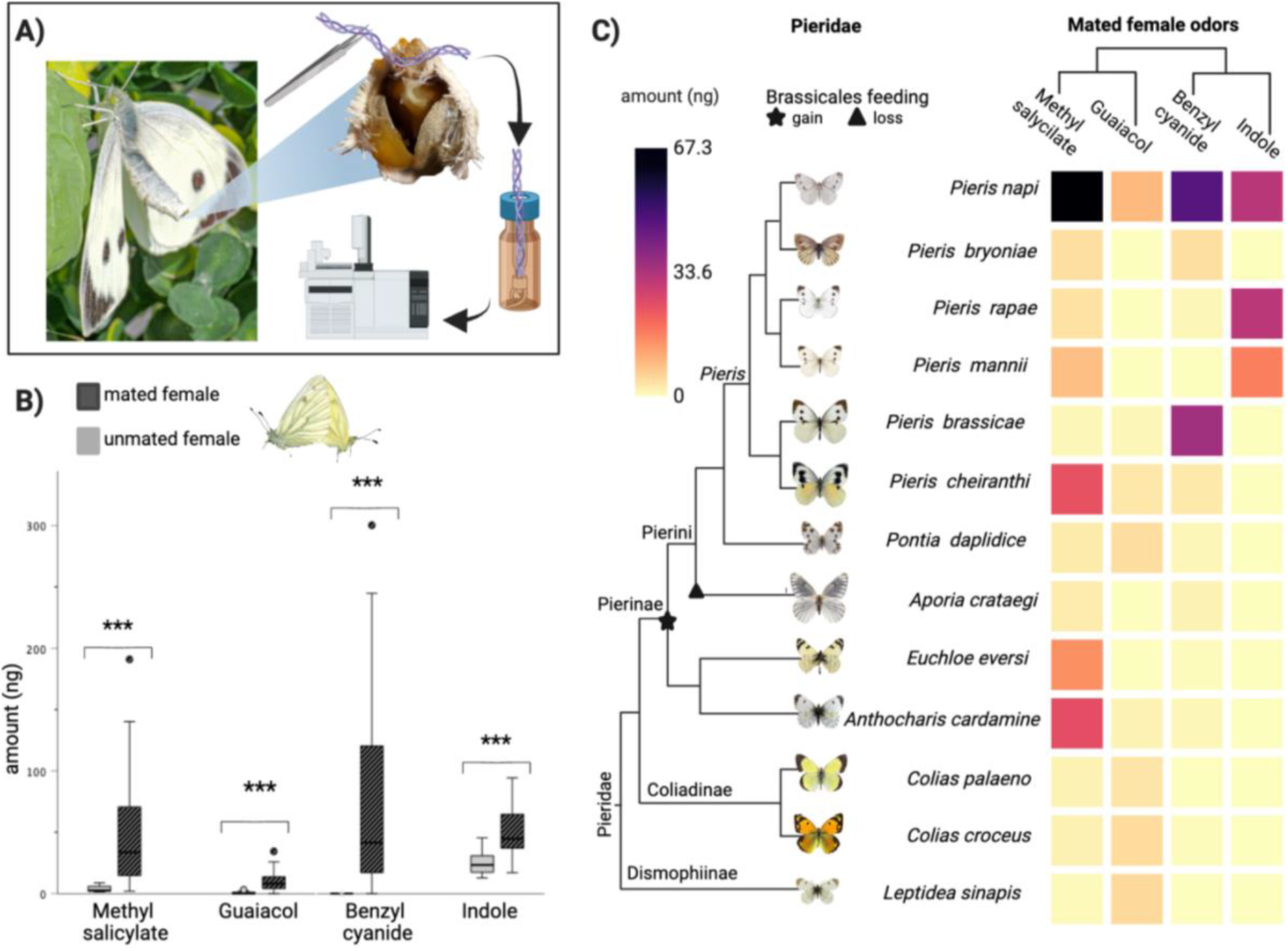
Post-mating odors are shared between Pieridae species. **(A)** Schematic of odor collection from wild-caught mated females using PDMS fibers applied to abdominal structures^42^. Inset: mate-refusal posture of a female (photo mate-refusal posture: Hans Smid, magnified female abdomen: Jonne Bonnet); collection with PDMS fibers; elution in hexane; analysis with gas chromatography. **(B)** Emission of four post-mating compounds (ng ± s.e.) in unmated and mated *Pieris napi* females from a Dutch population. Boxplots show data for 5-day-old females, either unmated or mated 24–48 hours prior. Asterisks indicate significance (***: p < 0.001; statistical details in supplementary methods and Table S2). **(C)** Emission profiles of the four components of post-mating odors, namely methyl salicylate, guaiacol, benzyl cyanide and indole across 14 wild-caught mated Pieridae species. Left: phylogeny (after Wiemers et al.^75^). Right: heatmaps show compound quantities (ng) relative to internal standard (pentadecane). Compounds are clustered by chemical distance. Statistical details in supplementary methods and Table S3.

Our analysis showed that all four compounds, including those previously considered possible biosynthetic by-products^14^, are robust and reliable indicators of mating in *Pieris napi* because they are present in much higher quantities in mated versus virgin females. Notably, our analysis used Dutch populations of *P. napi* and direct-contact volatile sampling, in contrast to earlier studies that relied on headspace collection. As headspace methods capture volatiles emitted during active mate refusal, while our method samples directly from the relevant tissue, both qualitative and quantitative differences may arise^42^. Manipulation of the butterflies during handling for collection may also enhance the release of certain compounds, as observed in other insects, such as locusts, which emit more benzyl cyanide when disturbed by predators^20^.

Across 14 wild-caught Pieridae species, these compounds were consistently present in mated females, with substantial overlap among species (Fig. 1C). This pattern contrasts with the high species-specificity typical of most lepidopteran sex pheromones and challenges previous assumptions of species-specific anti-aphrodisiacs^12^, suggesting that post-mating odor profiles are shaped by shared ecological factors rather than divergent sexual selection^39^. Methyl salicylate was the most widespread, detected in most species, consistent with its broad use as a chemical cue among Lepidoptera^44–46^. Indole showed the highest specificity, restricted to *P. napi, P. rapae*, and *P. mannii*. Benzyl cyanide was limited to Pierini, although this may reflect sampling bias; given that cyanogenesis occurs in other Pieridae and distantly related lepidopterans^47,48^, a broader evolutionary origin is likely. As expected for compounds derived from common metabolic precursors, such as phenylalanine, the synthesis of methyl salicylate and benzyl cyanide appears independent of host-plant chemistry^12^. Indole, synthesized from tryptophan^12^, represents a derived biosynthetic trait, as it appears restricted to a single clade within *Pieris*. While methyl salicylate alone has been shown to deter male *P. napi*, the behavioral functions of benzyl cyanide and indole in this species remain untested^12^.

Taken together, the widespread occurrence of these volatiles across species indicates that post-mating odors are environmentally ubiquitous signals, likely subject to selection pressures from natural enemies more so than conspecific mate recognition. While pheromone specificity between species often depends on differences in ratios and blends, not compound presence or absence^5^, high variability within species and populations would indicate that these odors are not reliable for species recognition. Furthermore, because these odors are emitted only after mating has already occurred, there is little reason to expect them to exhibit species specificity despite previous assumptions of stable differences between *Pieris* species. Unlike aphrodisiac pheromones, which can function in pre-mating species recognition to prevent hybridization^1,5,39^, species-specific post-mating odors would offer no such selective advantage.

### Intra-specific variation in post-mating odors corresponds with parasitoid pressure but not geographic distance, genetic distance, or mating rates

Marked population-level differences in post-mating odor emission across seven *Pieris napi* complex populations were stable over two years (Fig. 2A, B). Strikingly, populations with low or trace emissions (JUR, BRY, PYR) showed no evidence of egg parasitism despite the presence of *Trichogramma* parasitoids on sympatric lepidopterans (Fig. 2C, D, Supplement). In contrast, sites with high emissions (COB, WAG, STK, ABS) experienced frequent parasitism and produced distinct blends of methyl salicylate, benzyl cyanide, or indole, though rarely all three. Y-tube olfactometer assays confirmed that *T. cacoeciae*, the dominant phoretic parasitoid on *P. napi* and the only species tested that showed innate attraction to mated *P. napi* (Fig. 2F), was attracted to volatiles from mated females in high-emitting populations, but not to odors from low-emitting populations (Fig. 2G). These results demonstrate that post-mating odors in *P. napi* function as long-range attractants for phoretic parasitoids and that emission levels correspond with local parasitism risk.

**Figure 2.**
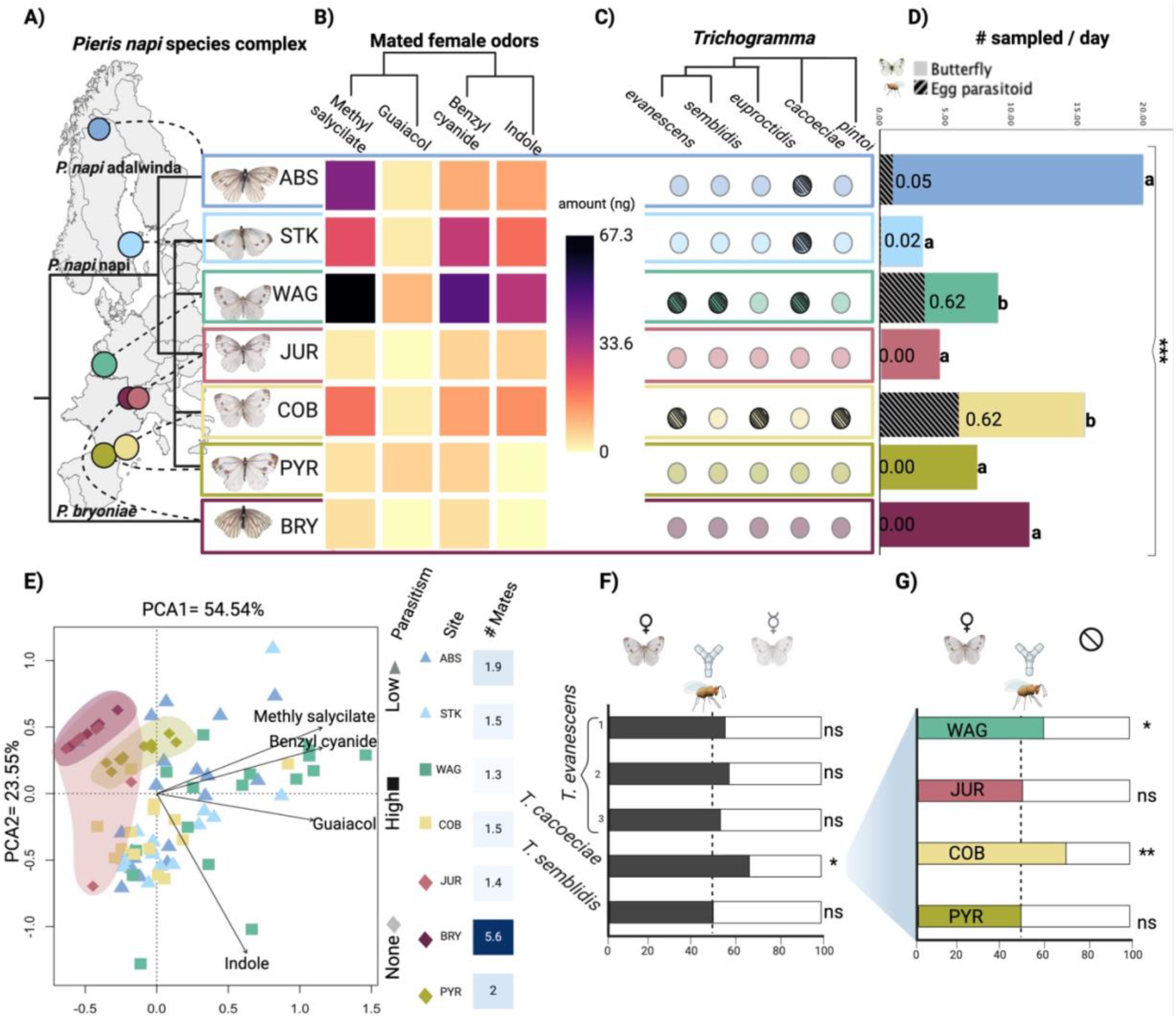
Intraspecific variation in *Pieris napi* post-mating odors is shaped by parasitoid pressure rather than geography or genetic distance. Asterisks indicate significance (*p < 0.05, **p < 0.01, ***p < 0.001, ns = p > 0.05). **(A)** Sampling locations and COI phylogeny of *P. napi* complex populations across a European latitudinal gradient sampled over two consecutive years (2022 & 2023). Circles on the map indicate collection site, details in supplementary methods. **(B)** Heatmap showing mean emission (ng) of four post-mating volatiles per population across two years, with compounds clustered by chemical distance. Sample sizes are in Table S4. **(C)** *Trichogramma* species detected in butterfly eggs and hitchhiking on butterflies by population; wasp phylogeny inferred from COI barcodes. Darkened circles indicate presence. **(D)** Daily capture rates of *P. napi* and *P. bryoniae* butterflies (color bars) and *Trichogramma* wasps (grey slashes) per site; bar labels indicate egg parasitism rates. Letters denote statistically distinct groups. Sample sizes in table S5. (E) Principal component analysis (PCA) of individual odor profiles by population (colors from panel A). Shapes indicate parasitism level: diamonds (none), triangles (low), squares (high). Blue boxes show the mean spermatophore number per population. Clusters JUR, BRY, and PYR were highlighted with hand drawn ellipses to encompass all points in their respective colors. **(F)** Y-tube olfactometer assays: proportion of naïve *Trichogramma* attracted to mated vs. unmated *P. napi* (WAG). Black bars indicate a preference for mated females. Statistical details in table S6. **(G)** Y-tube olfactometer assays, *T. cacoeciae* choice between clean air and mated females from four populations (colors from panel A); colored bars indicate butterfly choice. Statistical details in table S7.

Odor variation was not associated with geography, genetic structure, mating rates, or host-plant. This pattern held both in our own dataset and when compared to previously published findings on *P. napi*, which included more detailed genomic, host use, and mating analyses^37,49–51^. For example, genetically divergent *P. napi* adalwinda (ABS) and *P. napi* napi (STK, WAG, COB) exhibited similar blends, while distinct species *P. napi* and *P. bryoniae* from the same site (JUR and BRY) shared low emission levels (Fig. 2A, B). Only *P. bryoniae*, which emits little post-mating odor, showed high mating rates, while all other populations exhibited average remating frequencies regardless of odor emission (Fig. 2E). This suggests that post-mating volatiles are not primarily constrained by mating system or genetic drift, consistent with previous findings that divergence in *P. napi* and *Heliconius* polyandry does not correspond with differences in spermatophore derived compounds^34^.

Emissions of methyl salicylate and benzyl cyanide were strongly correlated and inversely related to indole, indicating a metabolic trade-off or alternative regulatory mechanisms (Fig. 2B, E). Methyl salicylate and benzyl cyanide derive from phenylalanine, while indole is synthesized from tryptophan^12^, representing alternative biosynthetic pathways. Methyl salicylate can also arise from benzyl cyanide, as caterpillars fed benzyl cyanide incorporate it into methyl salicylate, though *P. napi* males can also synthesize methyl salicylate *de novo* from precursors acquired by larvae and replenish adult stores by nectar feeding^14^. Notably, it has been found that indole does not attract other parasitoids^52^, possibly due to its toxicity to insects^53,54^, which may confer an ecological advantage and help maintain odor variation within populations^5,55^.

These data indicate that while anti-aphrodisiac use is widespread and conserved across Pieridae, its expression within *P. napi* is highly variable and ecologically contingent. In the absence of detectable effects on mating behavior, high levels of post-mating odor production likely incur ecological costs. Populations emitting low or no volatiles may escape detection, consistent with the idea that post-mating signal divergence is shaped by enemy-mediated selection rather than mate recognition. Alternatively, parasitoid preferences and cue use may themselves evolve in response to local odor availability, potentially driving reciprocal adaptation in this system. Time-calibrated analyses of biosynthetic and olfactory gene evolution in both butterflies and parasitoids could further illuminate the dynamics of this diffuse coevolutionary interaction^56^.

### Conserved cross-species and cross-sex olfaction of post-mating odors

Post-mating odors have traditionally been considered species-specific signals, with benzyl cyanide thought unique to *P. brassicae*^57^. However, given the widespread emission of methyl salicylate, benzyl cyanide, and indole across *Pieris* species (Fig. 1), we tested whether antennal sensitivity to these compounds is similarly conserved.

Using electroantennogram (EAG) recordings, we measured dose-response curves in males and females of four sympatric *Pieris* species (*P. brassicae, P. napi, P. rapae*, and *P. mannii*) from the Netherlands. Contrary to expectations of species- or sex-specific detection, all species and sexes exhibited remarkably consistent antennal responses to benzyl cyanide, methyl salicylate, and indole (Fig. 3A). Benzyl cyanide elicited the strongest responses, exceeding those to plant volatiles. In contrast, sensitivity to methyl salicylate and indole resembled responses to other common host-plant cues. This conserved detection likely reflects ancestral olfactory capabilities co-opted for post-mating signaling^58^, as these compounds also serve as oviposition cues in host-plants^18,59,60^. The similar sensitivity across species, including for *P. napi*, which was not previously known to use benzyl cyanide, challenges the notion of strict species-specificity in anti-aphrodisiac reception.

**Figure 3.**
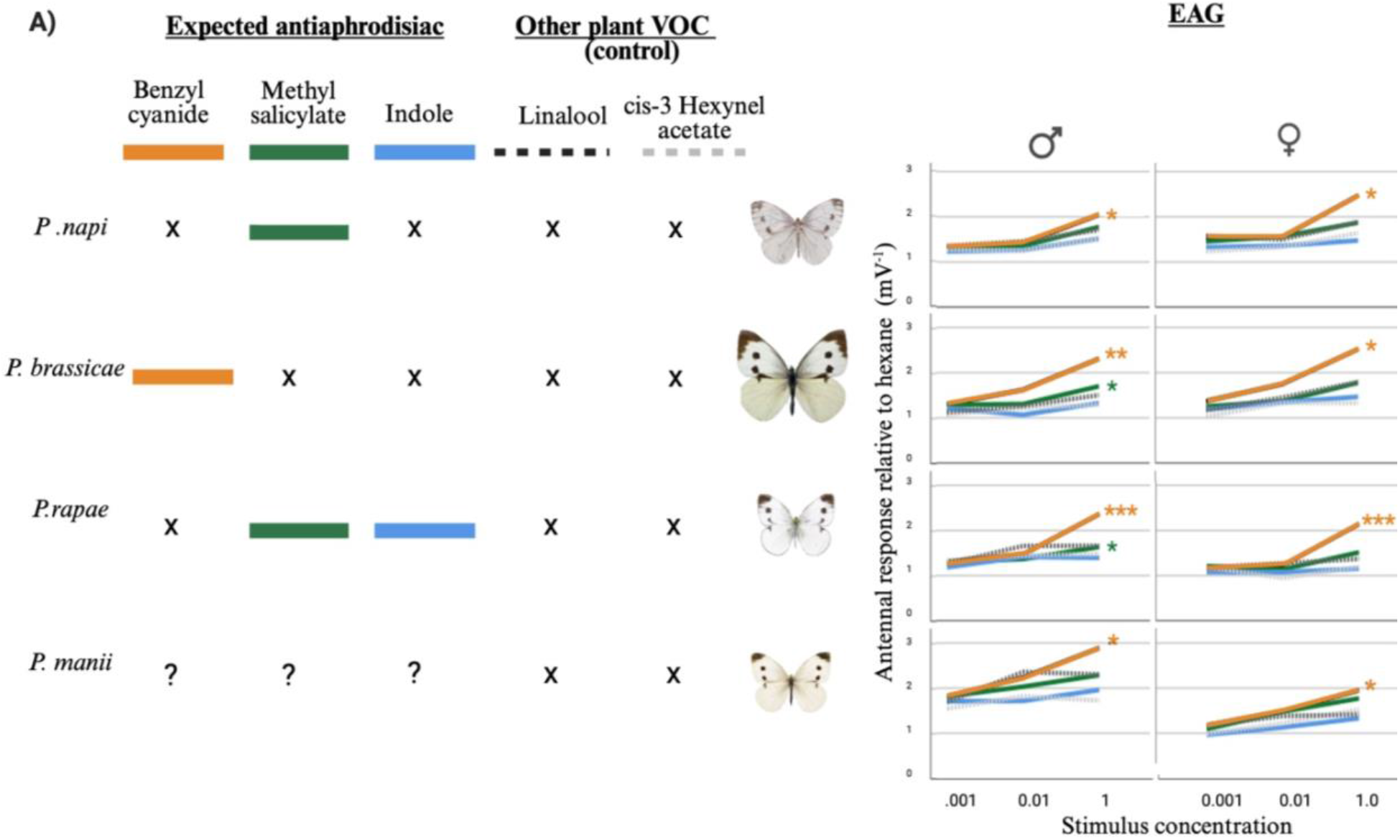
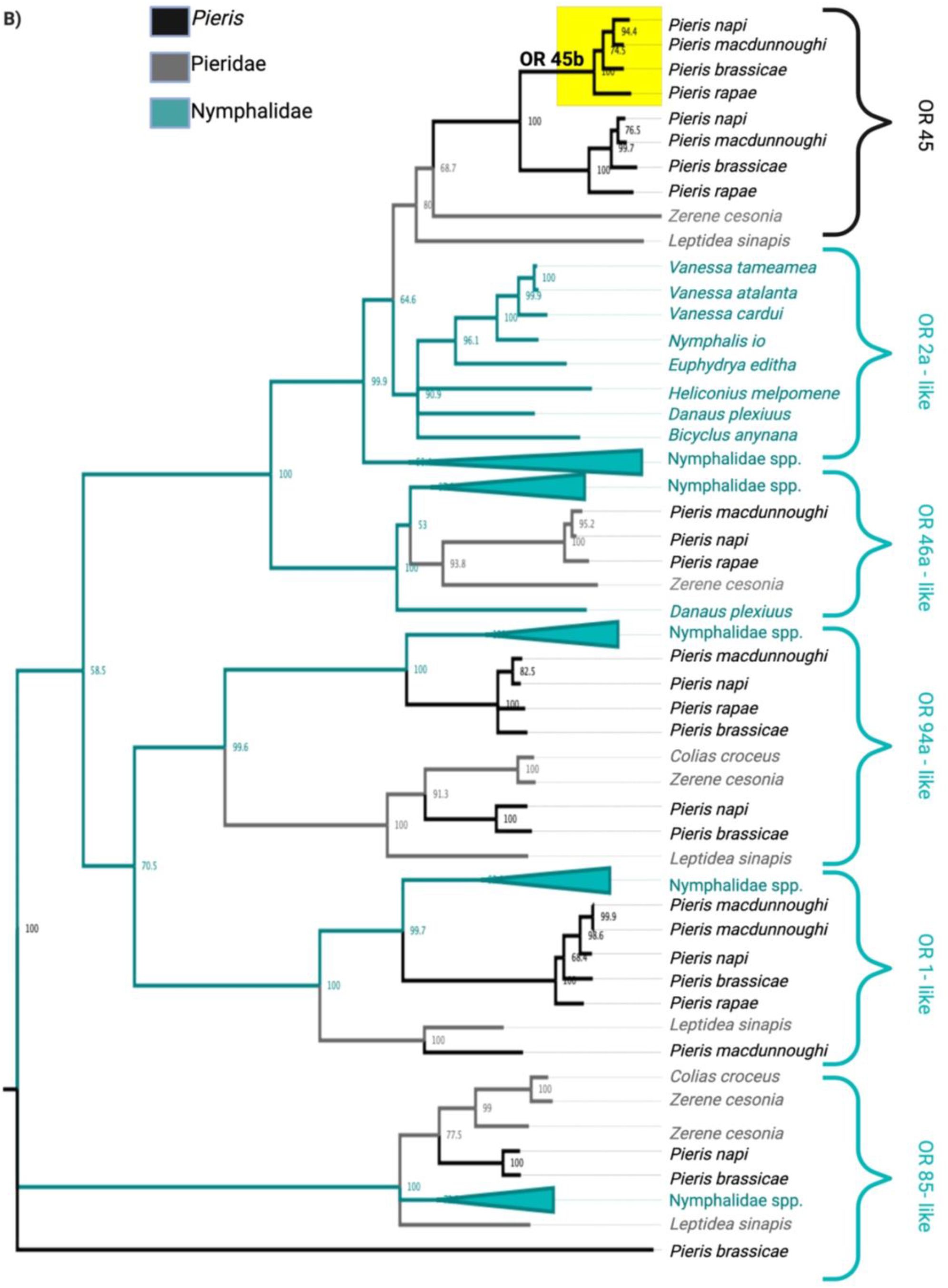
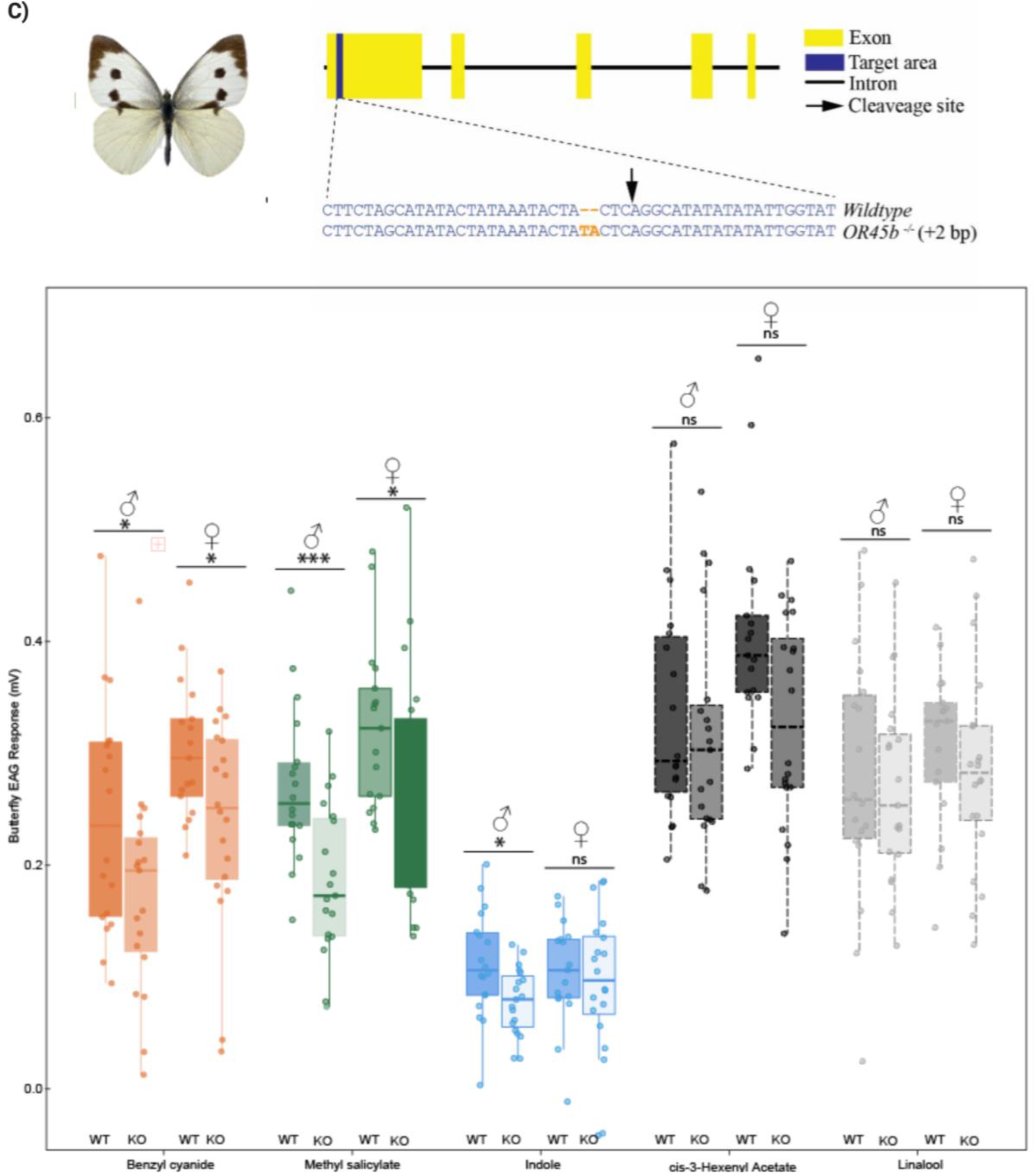
Interspecific variation and genetic basis of olfactory responses to post-mating odors.**(A)** Electroantennogram (EAG) dose-response curves of males and females from *P. brassicae, P. napi* (WAG), *P. rapae*, and *P. mannii* to conspecific and heterospecific anti-aphrodisiac compounds (benzyl cyanide, methyl salicylate, indole) and control plant volatiles (cis-3-hexenyl acetate, linalool). Colored lines indicate compound identity as indicated in the top legend; “X” marks compounds not known as anti-aphrodisiacs in that species. Responses are normalized to solvent control. Asterisks indicate significant effect of dose on response for each compound (*p < 0.05, **p < 0.01, ***p < 0.001, ns = p>0.05). Statistical details can be found in tables S8 and S9. **(B)** Maximum likelihood phylogeny of OR45b and homologs in Pieridae and Nymphalidae, rooted on *P. brassicae* ORCO, highlighting conservation of this receptor unique to *Pieris*.**(C)** Schematic of CRISPR/Cas9 knockout of OR45b in *P. brassicae* and EAG responses of wild-type (WT) and OR45b knockout (KO) butterflies to volatiles from panel A, demonstrating decreased post-mating odor detection in KO males and females. Asterisks indicate statistical significance: (*p < 0.05, **p < 0.01, ***p < 0.001, ns = p>0.05). Statistical details for these five compounds are found in table S9 and full EAG panel results are in Figure S3.

Comparative genomic and antennal transcriptome analyses^61^ indicate that *Pieris* chemoreception is conserved across species, with differences arising primarily from expression level variation. OR45b stands out as a *Pieris*-specific and conserved odor receptor in this genus (Fig. 3b). Knockout of OR45b indicates this receptor as a candidate partially mediating post-mating odor detection. CRISPR/Cas9 knockout of OR45b in *P. brassicae* significantly reduced antennal responses to these compounds, especially benzyl cyanide and methyl salicylate, without affecting responses to other plant volatiles (Fig. 3C), confirming its functional role.

The similar antennal sensitivity in males and females is notable. Females have been thought to lack control over anti-aphrodisiac emission^35^, with odor titers decreasing steadily with each emission event when the mate-refusal posture is adopted^15,62,63^. However, females’ ability to detect these compounds may serve to regulate remating timing and avoid autotoxicity from benzyl cyanide, which metabolizes to dangerous hydrogen cyanide^20^. Potential female self-regulation of anti-aphrodisiac levels warrants further investigation. In other Pieridae, such as wood whites (*Leptidea* spp.), only females can discriminate between con- and hetero-specific mates, which only differ in subtle amounts of chemical cues^64–66^. That male *P. napi* tailor ejaculate dependent on female anti-aphrodisiac emission^67^ may also provide a degree of protection from dangerous odor accumulation or emission.

### Functional consequences of post-mating odor perception

If post-mating odors are not essential for mating and increase parasitoid attraction, why do butterflies use them at all? To address this question, we examined the functional consequences of post-mating odor perception and emission in *P. brassicae*.

Disruption of the *Pieris*-specific odorant receptor OR45b^61^ in *P. brassicae* reduced antennal responses to post-mating odors (Fig 3C). OR45b knockout females laid significantly more eggs on previously infested host-plants. The knockout butterflies also showed reduced avoidance of oviposition-induced plant volatiles compared to wild-type females (Fig. 4A, B, supplement), both indicating impaired host discrimination. Methyl salicylate, benzyl cyanide, and indole are implicated in host-plant selection^16– 18,68^, and benzyl cyanide is deposited with *P. brassicae* eggs^27,69^. A reduced olfaction of these compounds likely impaired the ability of the knockout butterflies to detect heavily egg-laden plants, which would have been poor choices for oviposition because larval competition would be high. Knockout females were also more attractive to *Trichogramma* parasitoids than wild-type females (Fig. 4C). We expect that this altered parasitoid response is due to knockout females also emitting higher levels of benzyl cyanide due to a loss of feedback regulation, though this remains to be tested. If wild-type butterflies can reduce emissions when environmental titers of this risky compound exceed a certain threshold, e.g. by avoiding adopting the mate-refusal posture^63^, this would further suggest that post-mating odor detection enables females to regulate emissions, thereby optimizing both host selection and enemy avoidance. Future studies should investigate odor emission and adoption of the mate-refusal posture between these wild-type and mutant butterflies.

**Figure 4.**
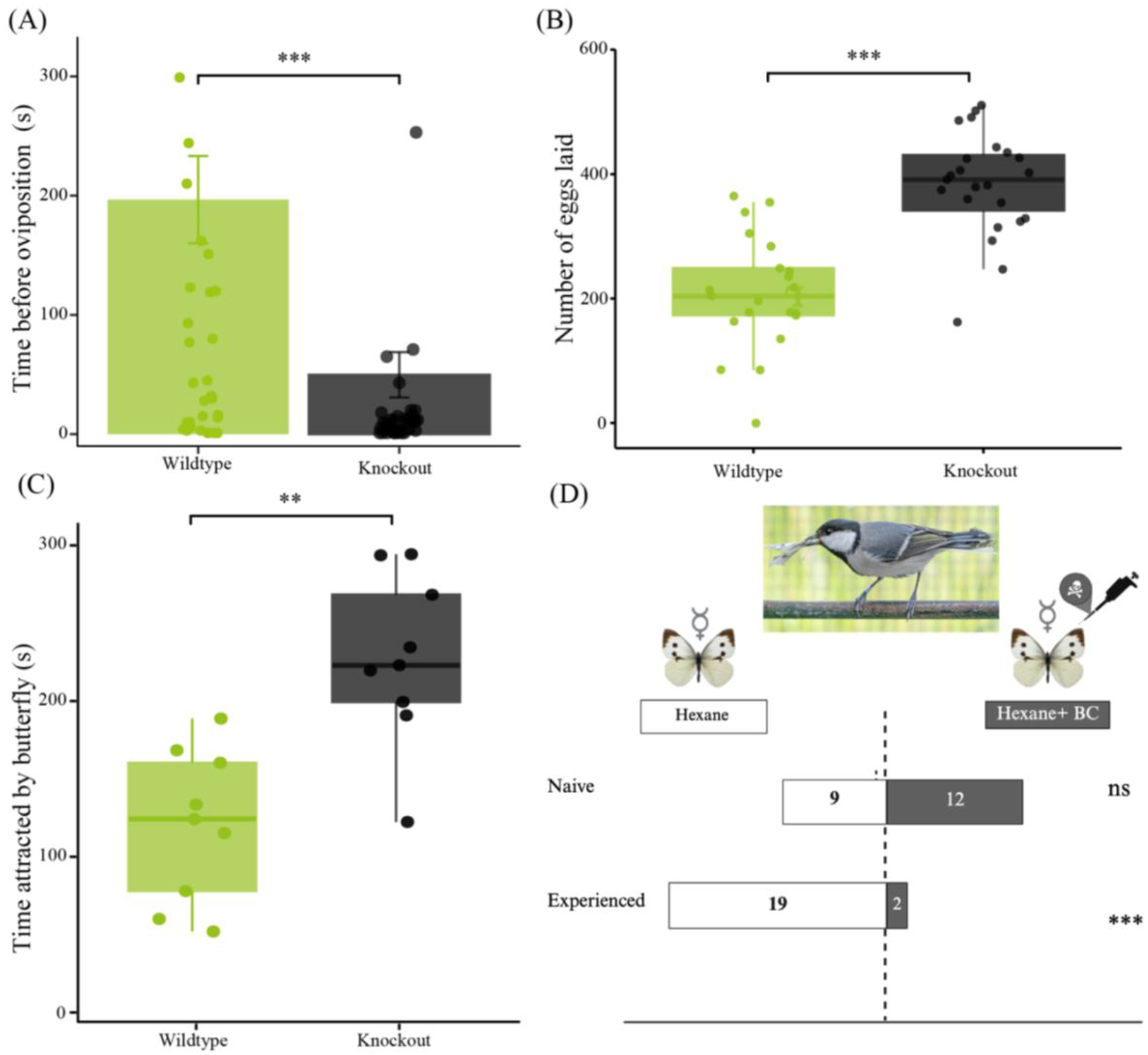
Functional consequences of post-mating odor perception and emission in *Pieris brassicae*. All statistical tests and sample sizes are provided in supplementary methods. Asterisks indicate significance: *p < 0.05, **p < 0.01, ***p < 0.001, ns = p > 0.05. **(A)** Time spent by wild-type (WT) and OR45b knockout (KO) females selecting host-plants. **(B)** Total number of eggs laid by WT and KO females on a single plant (not replaced) over one week. **(C)** Time spent by *Trichogramma* wasps above virgin and mated WT or KO females. **(D)** Naïve great tits (*Parus major*) given a choice between two virgin *P. brassicae*, one treated with 1–10 ng benzyl cyanide. The first attack choice was recorded after 10 minutes and repeated with the same birds in a second trial, now experienced. Statistical details in Table S10. Inset: photo of *P. major* eating *P. brassicae* by Hans Smid.

### Benzyl cyanide as an olfactory aposematic signal

Recent findings in locusts have shown that benzyl cyanide functions as an olfactory aposematic signal, warning predators of toxicity rather than serving strictly as an anti-aphrodisiac^20^. In our system, where butterflies emit even lower amounts of benzyl cyanide, this hypothesis is supported by behavioral assays with great tits (*Parus major*): while naïve birds attacked benzyl cyanide-treated and untreated butterflies equally, experienced birds reliably avoided treated individuals (Fig. 4D). This learned aversion enables them to avoid ingesting benzyl cyanide, which is converted to hydrogen cyanide upon attack, poisoning predators^20,47,48^.

Classic experiments showed that mice injected with homogenates of *P. brassicae* and *P. napi* died rapidly, even when the butterflies were reared on glucosinolate-free diets, whereas *P. rapae* homogenates were non-lethal^70^. Consistent with the *de novo* synthesis of post-mating odors^14^, toxicity was maintained even when larvae were reared on artificial diets without glucosinolates^70^. While this finding puzzled the researchers, the endogenous nature of benzyl cyanide in *Pieris* explains toxicity without dependence on plant-derived toxin sequestration. This finding also suggests that *P. rapae* may benefit from mimicry, gaining protection from predators by resembling the more toxic congeners without incurring the physiological costs of compartmentalizing benzyl cyanide itself.

Under high parasitoid pressure, such mimicry may allow species to retain anti-predator benefits while reducing the risk of parasitoid attraction, either by emitting lower levels of benzyl cyanide (and methyl salicylate) or by shifting to alternative compounds such as indole. *P. brassicae*, with its high benzyl cyanide concentrations, is among the most unpalatable members of a putative mimicry ring, supported by its distinctive black-and-white warning coloration^70,71^. However, prior studies on mimicry and aposematism in cabbage whites often failed to account for sex or mating status, typically using dead models or not specifically including mated females that emit warning odors when alive, which may explain inconsistent findings regarding palatability^70–73^. Our results suggest that the full ecological and evolutionary significance of these chemical defenses may only be revealed when considering live, mated females in the mate-refusal posture.

### Synthesis and Conclusions

Our findings overturn the long-standing view that post-mating odors in *Pieris* function and evolve primarily as sex- and species-specific anti-aphrodisiac pheromones^14,15^. Instead, we reveal that these compounds, once thought to be private sexual signals, exhibit unexpectedly low interspecific variation in both emission and detection. These so-called “pheromones” are, in fact, ubiquitous volatiles, easily intercepted by a broad range of organisms. Our results show that both males and females of multiple *Pieris* species detect and respond to these compounds, challenging the idea of strict species or sex-specificity and suggesting evolutionary conservation of the underlying sensory pathways.

Meanwhile, these odors exhibit high intraspecific diversity. This diversity does not track drift or mating behavior but instead aligns with ecological pressures: populations emitting more odors face higher parasitism, while predator interactions and host-plant volatiles further shape odor profiles. Whether this variation reflects underlying genetic divergence or phenotypic plasticity remains unresolved, but the latter may offer a particularly adaptive response to shifting ecological pressures.

Why then do butterflies persist in using such broadly available cues, despite the clear costs of parasitoid attraction? Our data support a new framework: these volatiles act as multifunctional infochemicals shaped by balancing trade-offs of ecological selection pressures. Detection of post-mating odors aids females in host-plant selection and may mediate feedback regulation of emission, allowing individuals to optimize the trade-off between reproductive signaling and enemy avoidance. Notably, benzyl cyanide also serves as a deterrent against great tits, indicating protection against vertebrate predators. This dual role is also observed in other insects, such as locusts, where benzyl cyanide serves as an aposematic signal^20^, which may explain its evolutionary persistence in *Pieris* despite the risks of parasitoid eavesdropping.

Our results further suggest that the evolutionary interplay between toxicity, mimicry, and chemical signaling may shape the diversification of post-mating odors in cabbage whites. Species like *P. rapae*, which emit no benzyl cyanide, may benefit from mimicry, gaining predator protection by resembling more toxic relatives while minimizing the costs of parasitoid attraction. In contrast, high-emitting species such as *P. brassicae* occupy the core of a mimicry ring, supported by both chemical and visual aposematism.

More broadly, our study calls for a rethinking of insect pheromones that co-opt common plant volatiles. Rather than functioning as isolated sexual signals, these compounds operate at the intersection of mating, oviposition, predation, and ecological community dynamics. In *Pieris*, post-mating odors reflect not only reproductive status but also the evolutionary entanglement of sexual selection, host adaptation, and enemy-mediated selection. This multifunctionality may be the rule rather than the exception for broadly detectable chemical signals in complex ecological networks.

## Methods

Complete protocols, reagents, and additional analyses are provided in the Supplementary Methods.

### Study system and sampling

We studied seven populations of the *Pieris napi* species complex and 12 other *Pieris* species across a European latitudinal gradient in 2022 and 2023. Adult butterflies were collected in the field and reared under controlled conditions. Individuals were captured in containers to check for phoretic *Trichogramma* and parasitized eggs were collected from host-plants to assess parasitism rates and rear parasitoids for identification. Mating frequency was estimated by counting spermatophores in dissected wild females. Sampling details are provided in Supplementary Table S1.

### Insect and bird rearing

Butterflies were reared on *Brassica nigra* under standardized greenhouse conditions (25 °C, 30% RH, 18:6 h light:dark). *Trichogramma* were maintained on *Ephestia kuehniella* eggs. Birds (Parus major) were reared at the Netherlands Institute of Ecology (NIOO-KNAW) under ethical approval.

### Insect identification and phylogenies

Species identification for *P. napi* species complex and parasitoids was based on CO1 barcodes, sequenced following Chelex DNA extraction and PCR with Folmer primers^74^. Sequences were aligned using MAFFT, and maximum likelihood trees were constructed in Geneious Prime (100 bootstraps, default settings).

### Chemical analyses

Post-mating odors were collected from live females who were found ovipositing on plants in the field using direct sampling with PDMS fibers^42^ by rubbing fibers against the abdomens of females found laying eggs. Samples were eluted in hexane and analyzed via gas chromatography using synthetic standards. Emission rates of benzyl cyanide, methyl salicylate, indole, and guaiacol were quantified relative to internal pentadecane standards. Virgin vs. mated *P. napi* were compared using lab-reared individuals. Chemical distances for clustering were calculated with ChemmineTools.

### Electrophysiology

Electroantennogram (EAG) recordings were performed on excised antennae from three-day-old butterflies (n = 15–20 per group) exposed to synthetic odorants. Responses of four *Pieris* species were measured against three anti-aphrodisiac compounds and two control volatiles. Dose-response curves were generated using three dilutions. For *P. brassicae*, EAGs were recorded for wild-type and OR45b knockouts using 47 odorants (Fig. S3).

### Genetic and phylogenetic analyses of odorant receptor

Odorant receptor gene candidates were identified by comparative genomic analysis using protein BLAST on NCBI against all Pieridae and Nymphalidae. Maximum likelihood phylogenies of odorant receptor genes were constructed using protein alignments. For genetic analyses, CRISPR/Cas9 was used to generate OR45b knockout lines in *P. brassicae*. Guide RNAs targeting the first exon of OR45b were designed using Geneious Prime and synthesized via in vitro transcription. Newly laid eggs were microinjected with sgRNA/Cas9 complexes, and resulting caterpillars were reared to adulthood. Knockout individuals were identified by PCR and sequencing of leg tissue, with non-edited siblings serving as wild-type controls.

### Behavioral assays

Oviposition assays involved greenhouse trials with wild-type and knockout females exposed to host-plants and allowed to mate freely. Mating rates were determined (Fig. S4), and egg-laying was monitored daily for seven days (Fig. S4). Crosses among wild-type and knockout individuals tested for sex-specific effects (Fig. S4). *Trichogramma* responses were assessed using olfactometers: static assays compared *P. brassicae* knockout vs. wild-type, and Y-tube assays tested the attraction of naïve wasps to reared *P. napi* odors. Bird assays used naïve and experienced great tits to test preference between benzyl cyanide-treated and control butterflies (1–10 ng). Predator choice was recorded over sequential morning and afternoon trials.

### Statistical analysis

Statistical analyses, sample sizes, and significance thresholds are detailed in the Supplementary Methods and Tables S2–S4. Analyses were conducted in R and SPSS.

## Acknowledgments

For additional help with data collection, we thank Frank Rooijakker, Severin te Lindert, Jade Kandelaar, Samu Oostenhoek, and Teun de Jong. For support with lab and field work, we thank Constanti Stefanescu, Carl Gotthard, Joost van de Heuvel, Gabriel Joachim, Estrella Hernandez-Suarez, Jorge Alfredo Reyes-Betancort, and Eileen Bader. For discussions which helped with conceptualization of this study, we thank Heiko Vogel, Krister Wiklund, Chris Wheat, and Magne Friberg. For rearing insects, we thank Jordy Litjens, Patrick Verbaarschot, Violet van Oosten, and Gabriella Kiss-Bukovinsky. For photography, we thank Hans Smid. For funding, we thank the Dutch Research Council (NWO open-competition grant (OCENW.M20.027) and the KNAW academy ecology fund (Grant KNAWWF/747/ECO2021-11).

