## Supplement for "Enemies, more than sex, shape butterfly post-mating odor evolution"

### Supplement for

### Enemies, not sex, as a driving force of butterfly post-mating odor evolution

#### Methods

##### S1 Field surveys

***S1.1 Sites***

To assess variation in post-mating odors of *Pieris napi* and other Pieridae and their inter-actions with *Trichogramma* in geographically distant locations, we sampled six different populations of *P. napi*, its subspecies, and sister species across Europe. Sites were selected based on citizen science data indicating that *Pieris napi* and its close relatives were abun-dant. Sources included the Catalonian butterfly monitoring scheme (CatalanBMS.org), Obervations.org, INaturalist.com, and Artportalen.SE. At each site we also collected other Pieridae species we encountered.

Samples were collected in 2022 and 2023 from May to July in natural and cultivated sites. Each sampling site consisted of 2-5 transects within a area with a 50km radius (figure S1). Each transect was visited at least twice per sampling year, with sampling at least once in the morning and once in the afternoon. The coordinates listed below are the CenterPoint of each sampling area.

The Abisko (**ABS**) site is located in Northern Sweden (68° 2’21.95”N, 19°26’46.16”E). The sites around Abisko were characterised by big patches of flowering host plants like Bar-barea vulgaris and various *Arabidobsis* species. From the *Pieris* genus only the subspecies *Pieris napi* adelwinda Fruhstorfer was present, which occurs in one generation per year. At this site, we also collected *Colias palaeno*.

The Stockholm (**STK**) site is located in Southern Sweden (59°12’3.61”N, 17°45’21.31”E). We collected *Pieris napi* napi in city parks of Stockholm and Tovetorp biological station. In the city, *Pieris* were found mainly on *Bunias orientalis* and *Allaria petiolata*. Around Tovetorp, the landscape was characterised by forest interspersed with flowering meadows. *P. rapae* and *P. brassicae* were also present, and the three *Pieris* species generally have two genera-tions per year and are generally found on sparsely occurring *Barbarea vulgaris*. At this site we also collected *Leptidea sinapis*.

The Wageningen (**WAG**) site is located in the center of the Netherlands (51°54′41.53″N, 5°39′14.48′E). Around Wageningen the field sites were characterized by cultivated pastures with crops (many varieties of *Brassicae oleracea*) and wild-flower strips with *Raphanus raphanistrum*, *Sinapis arvensis* , and *Brassica nigra*. Here, three to four generations of *P. napi* napi, *P. rapae*, *P. brassicae*, and *Pieris mannii* co-occur and were all collected. We also collected *Anthocharis cardamine*, which were using *Cardamine pratensis*.

The Jura (**JUR**) site is located in the Jura Mountains on the French side of the border between France and Switzerland. (46°31′2.74″N, 6° 4′55.76″E). The field sites around Belfontaine, in the Jura mountains were characterised by meadows with flowering plants. At this site, both *P. napi* napi and *Pieris bryoniae* Ochsenheimer (Lepidoptera:Pieridae) were abundant and found using the host plant *Cardamine heptaphylla*. *P. bryoniae* (**BRY**) is a very close relative of P. *napi*, and in some sites where they co-occur they are known to hybridize. We also collected *Aporia crataegi* at this site.

The Costa Brava (**COB**) site is located in the north-eastern coast of Catalonia, Spain (42°13′22.91″N, 3° 0′50.31″E). The field sites in this area were characterised by grazing pastures interspersed where *P. napi*, *P. rapae*, and *P. brassicae* were all abundant and generally have four to five generations per year. *P. napi* was found near streams at the shaded edges of fields using the plant *Lepidium draba* and *Brassica nigra*. At this site, we also collected *Colias croceus*.

The Pyrenees (**PYR**) site is located on the Spanish side of the Pyrenees mountains, bordering France and (42°23′46.59″N, 1° 3′49.78″E). We collected mainly around Gerri de la sal, a dry mountainous area characterized by salt flats and high biodiversity. *Pieris napi* napi were less common, and found alongside streams on infrequent host plants such as *Allaria petiolata*. Here, we also collected *Pieris rapae*, *Leptidea sinapis* and *Pontia daplidice*.

Apart from our sampling of *Pieris napi* populations, further sampling of *Pieris chieranthi*, *Euchloe eversi*, and *Pieris rapae* was conducted in Tenerife and La Palma, in the Canary Islands, at 13 sampling sites selected in coordination with the ICAE Tenerife Agricultural Center. *Pieris napi* and *Pieris brassicae* do not occur there. *Pieris chieranthi* were collected in La Palma.

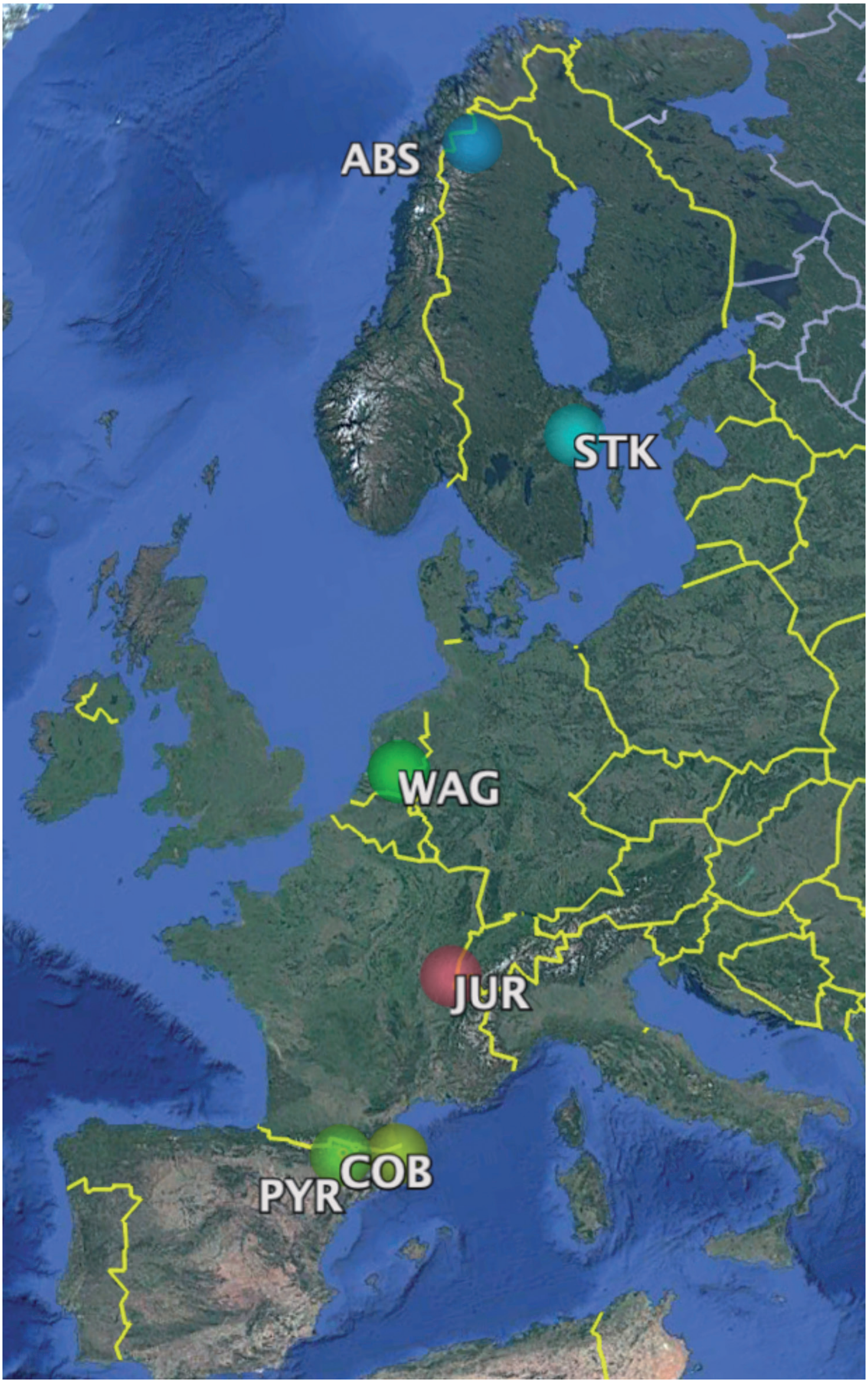


**Figure S1)** Sampling sites across Western Europe. The overlaid circles represent the sampling areas, with a radius of 50 km. ABS= Abisko, STK = Stockholm, COB= Costa Brava, WAG = Wageningen, JUR= Jura mountains, COB= Costa Brava, and PYR = Pyrenees mountains.

##### *S1.2 Phoresy and parasitism rates*

Sample sizes for butterflies and wasps per location can be found in table S5. Eggs were collected from hostplants at the six different locations. Eggs were checked for signs of parasitism in the field. When eggs are parasitized they turn black within approximately 5 days since parasitiation. Eggs that still appeared yellow (unparasitized) were collected in plastic petri dishes with moistened filter papers and observed until either turning black or caterpillars hatched, at which point they were recorded as parasitized or unparasitized. Caterpillars that hatched from unparasitized eggs were put back in the environment of collection. Some caterpillars were kept for starting a rearing of the population when back in Wageningen.

Parasitized eggs were isolated in small tubes to allow capture of hatching of the wasps. Empty eggshells were stored in 96% ethanol for DNA analyses to identify the host species when both *P. rapae* and *P. napi* were present, as they use similar host plants and their eggs cannot be morphologically distinguished (Chapter 4). However, identification of eggs to species label was not always possible and parasitism rates are calculated as the proportion of parasitized versus unparasitized eggs for all solitary Pieris eggs.

Emerged wasps were relocated to a plastic tube and provided with a droplet of biological honey. The sex of parasitoids was established by looking at the antennae. When a female wasp was present, the wasps were provided with reared on *Ephestia kuehniella* Zeller (Lepidoptera: Pyralidae) obtained from Koppert Biological Control (The Netherlands) to start a rearing. Dead specimens were stored in 96% ethanol for DNA analyses.

Butterflies were caught to survey for phoretic *Trichogramma* following the method described by Fatouros & Huigens (2012), using a butterfly net (Ø 40cm, Vermandel, The Netherlands). At the bottom of the net, a plastic container (Volume 250ml) was secured with a rubber band. In one fluid sweep of the butterfly net, the butterflies were caught in the air without touching the vegetation surrounding, to minimize bycatch. After the butterfly entered the container, the container was closed immediately with a lid to maximize the chance of capturing all possible hitchhikers present on the butterfly. After every catch, the container was replaced by a new empty container. After every three catches, a control sweep was performed, in which the same sweeping motion was used. For the control catches, instead of catching a butterfly, only air was sampled, and the container was checked for presence of bycatch. The control sweeps did not yield any *Trichogramma* wasps.

Phoresy rate was determined in the field by catching *Pieris napi* and checking them for possible hitchhikers. The butterflies were thoroughly checked, two times with a ±30 minutes interval. Special attention was given to the head, thorax and wings of the butterfly.

When a possible hitchhiker was spotted the container with the butterfly was retrieved to the lab. The containers were put in the fridge at 4°C for 5-10 minutes to slow down the butterfly and possible hitchhikers. Possible hitchhikers were separated from the butterfly and put in a plastic test tube with a drop of biological honey and eggs to parasitize. Depending on the sex of the butterfly, it was released (male) or kept for pheromone collection (female). Possible hitchhikers were reared and brought back to Wageningen. Laboratory populations were established and reared on *Ephestia kuehniella* Zeller (Lepidoptera: Pyralidae) eggs. Dead specimens were stored in 96% ethanol for DNA analyses.

##### *S1.3 Parasitoid identification*

Wasps were dried at 30 °C to minimize the amount of ethanol present in the samples. Sterile glass pestles for grinding samples were made by melting the tip of glass Pasteur pipettes into a rounded ball, with a Bunsen burner. Single *Trichogramma* wasps were grinded with a sterile glass pestle in a 0.5 ml Eppendorf tube. For each sample a new sterile pestle was used. 50 µl of 5% Chelex X-100 and 4 µl Proteinase K (20 mg/ ml) was added to each sample using a pipette and centrifuged to spin all content to the bottom of the Eppendorf tube. Samples were incubated for a minimum of 6 hours at 56°C. Samples were centrifuged for 1 minute and subsequently heated for 10 minutes at 95°C to inactivate Proteinase K. Finally, samples were spun down, and DNA extract was used for PCR. A polymerase chain reaction to amplify the Folmer barcode region (Folmer et al., 1994)[79] was performed using forward primer LCO1490: 5'-GGTCAACAAATCATAAA-GATATTGG-3' and reverse HCO2198: TAAACTTCAGGGTGACCAAAAAATCA-3'. The reaction was conducted in a total volume of 25 µl containing 20.5 µl of PCR Master Mix (12.5 µl 2X mix Phusion flash, 1.0 µl 10 mM forward primer, 1.0 µl 10 mM reverse primer, 6.0 µl H2O) and 4.5 µl of DNA template. The PCR cycling program consisted of a starting cycle of 10 sec at 98°C followed by 30 cycles of denaturation for 5 seconds at 98°C, annealing for 10 seconds at 58°C and polymerisation for 30 seconds at 72°C with a final elongation cycle of 60 sec at 72°C after the last cycle. Amplicons were run on a 1.5% agarose gel in 0.5 TAE at 100 V for approximately 30 minutes. The gel was stained with ethidium bromide (0.5 µl EtBr per 100 ml gel) and visualized with UV light. Trichogramma amplicons detected by electrophoresis with the expected band size (550 bp in size) were prepared for sequencing bidirectional Sanger sequencing by Eurofins genomics. Sequences were prepared using Geneious prime 2023.0.4. De novo assembly of forward and reverse reads, trimming, and generation of consensus (error probability limit 0.05) were performed with default settings. A consensus sequence was made using default settings for all sequences. Identification of the wasps was achieved by comparing the obtained FASTA-sequences with the Hymenoptera collection (nr/nt) database using BLASTN (NCBI interface). The identification of the FASTA-sequences was based on the E-value (>0.01), Percentage Identity (>95) and Max score. A phylogenetic tree was built by performing a multiple alignment of obtained *Trichogramma* sequences on Geneious prime 2023.0.4. Alignment was made with MAFFT (Katoh et al., 2009) using default settings. A maximum likelihood tree was generated using PHYML with 100 bootstraps. Geneious consensus tree builder was used to make a consensus tree.

#### S2 Rearing

***S2.1*** *Pieris napi*

Larvae of *P. napi* collected from eggs laid by 5- 10 wild butterflies during field work from May-July, 2022 and 2023 were reared at the greenhouses of Unifarm, Wageningen University & Research. Rearings were maintained under greenhouse conditions in ventilated nylon cages (adults: 120 cm3, larvae: 60 cm3) at 21°C, 60% relative humidity and a 18:6 h light:dark photoperiod. Adult butterflies were provided with a cotton wad soaked in 10% honey water solution and a *Brassica nigra* plant for egg-deposition. Every second day the cotton wad was refreshed and eggs were collected to maintain the population. Once populations were established with approximately 30 adults at a time. Approximately 100 eggs were collected every other week. Larvae were reared on *B. nigra* plants.

***S2.2*** *Pieris brassicae*

Seeds of cabbage plants (*Brassica oleracea* var. *gemmifera* cv. Cyrus; Brussels sprouts) were planted in individual pots and grown for four weeks after germinating. The cabbage plants were cultivated under glasshouse conditions with 22 ± 3 °C, RH 50-80 % and 16 h light: 8 h dark cycles. *Pieris brassicae* (Lepidoptera: Pieridae) caterpillars and butterflies were taken from the laboratory colony maintained at the Laboratory of Entomology, Wageningen University & Research, The Netherlands. The caterpillars were reared on four-week-old *Brassica oleracea* var. *gemmifera* cv. Cyrus, Brussels sprouts plants at 22 ± 3 °C, RH 60-80 % with a 14h light : 10h dark photoperiod until pupae. The male and female butterflies were fed with 5 % sugar water under the same environmental conditions as the caterpillars. The genetically modified insects are reared in a glasshouse compartment at Unifarm, Wageningen University & Research at 22 ± 3 °C, RH 60-80 % with a 14h light : 10h dark photoperiod. Mutant insects were supplied with the same food as wildtypes.

***S2.3 Wasps***

*Trichogramma* wasps collected during fieldwork from May-July, 2022, were reared under laboratory conditions in glass tubes (Ø 1cm) at 18°C, 50-70% relative humidity and a 16:8 h light:dark photoperiod at the Laboratory of Genetics of Wageningen University & Research. Adults were reared on egg-cards ( 50 x 150mm) made from *Ephestia kuehniella* eggs sourced from Koppert in The Netherlands secured to the cards with double-sided tape. The wasps were provided with a small droplet of organic honey. After emergence, male and female parasitoids were kept together for mating. Fresh egg cards were given every three weeks. For the experiments, females were isolated in glass tubes before being tested.

For bioassays with mutant butterflies, the egg parasitoid species used in this experiment was *Trichogramma evanescens* (strain 69_eng_20) (Hymenoptera: Trichogrammatidae). This iso-female line was reared from a single female wasp collected in Wageningen, The Netherlands and has since been as described above.

##### *S2.4 Birds*

For this study we will use great tits (*Parus major*) (n=17) housed at the Animal Ecology laboratory at the Netherlands Institute for Ecology (NIOO-KNAW), Wageningen. The birds are a F2 population and were born and raised in captivity. The birds originated from a population of urban birds. They were individually housed in cages of 90 x 50 x 30 cm under a photoperiod of 14:10 L:D and were fed on a diet of mealworms, songbird seed mix and apple pieces. Drinking water and bathing trays were freely available. Five sides of the cage are covered so the influence of external disturbances is as low as possible. The birds are used by esearchers and have been used in exploratory and behavioural studies before. They did not have any prior experience with Pierid butterflies or with benzyl cyanide.

#### S3 Odor samples

##### *S3.1 Collection in the field*

Per population 13-27 females of *P. napi* and per population and 2-10 individuals of other Pierid species were collected for pheromone collection (described in S1.1). To ensure we collected odors from mated females, we sampled from wild females that we witnessed laying eggs on host plants.

To determine the composition of post-mating odors, we collected odor samples with PDMS fiber rubs. A study by Lievers & Groot (2016) showed that pheromone composition collected with PDMS rubs closely resembles the pheromone composition collected with solid-phase micro-extraction (SPME), used by most previous studies into AA of *Pieris* (Lievers & Groot, 2016). All tools used during pheromone collection were thoroughly cleaned before and between each sample collection in ethanol and hexane.

Pheromone rubs were collected following the method described by Lievers & Groot. For the fiber rub method, fused silica optical fibers coated with a 100 µm PDMS layer (Polymicro Technologies Inc., Phoenix, AZ, USA) were cut into 20 mm pieces. The fibers were washed by gently shaking them in a bottle with hexane. Each fiber was subsequently wrapped air-tight in aluminum foil and stored in an aluminum foil envelope until use in the field. Pheromones were collected from female butterflies 2-6 hours after collection in the field. Pseudo mate-refusal posture was achieved by gently squeezing the abdomen of the butterfly between thumb and index finger, whereby the abdominal structures exposed during the mate-refusal posture were also exposed during this handling. With clean forceps, the fiber was rubbed over the abdominal structures, extruded by the female, for ±3 minutes. Each fiber was subsequently placed in a 0.05 mL clear glass micro-insert in a 2 mL clear glass crimp-cap vial. Odor collection vials consisted of inserts that were placed in a spring (36 x 5 mm) in a 2 mL clear glass crimp-cap vial (32 x 12 mm) and capped with a crimp cap with a PTFE (tetrafluoroethylene) septum. The vials were pre-filled with 50 ul of hexane and 200 ng of pentadecane as an internal standard. Fibers were then rinsed in the vials by gently tilting the crimp cap vials a couple of times. After 30-60 minutes, fibers were rinsed again and removed from the insert and sealed vials were stored at -20°C until further analysis.

##### *S3.2 Mated versus unmated* P. napi

To assess which of the pheromones are transferred by the males to the females, we compared the pheromones of virgin and mated females from the greenhouse rearing of the Wageningen population. To achieve this, we sampled 10 mated females and 20 virgin females per population following the PDMS fibre rub method described above. Virgin females were obtained by separating the butterflies by sex as pupae. After the emergence of the butterflies, 10 females were kept separated to assure virginity, the rest were combined with the male butterflies for mating. Butterflies were observed and all mating couples were directly isolated in separate nylon cages (15 $cm^3$). When mating was final-ized and the pairs separated, females were isolated in individual containers until odor collection to avoid them emitting odors when adopting mate refusal postures, which also occurs in response to other female butterflies, thereby keeping them from reducing titers. PDMS rubs were taken 36 hrs after mating. At the time of sampling, butterflies of
5-8 days old.

##### *S3.3. Odor analysis*

Pheromones were analyzed following the method described by Lievers & Groot, 2016 . For pheromone analysis, the volume of all extracts was reduced to 1–2 µl under a gentle stream of nitrogen. To prevent evaporation after this, the 2 µl samples were taken up together with 1–2 µl octane (Anhydrous 99+%) with a 10 µl glass syringe (701SN needle; Hamilton). The total volume of 2–4 µl was placed in a 50 µl glass insert, which was placed in a metal spring (35x5mm) within a 2 ml glass crimp vial, capped with an 11 mm aluminum crimp cap and a PTFE (tetrafluoroethylene) septum.

The pheromone extracts were injected with an Agilent 7693A Automatic Liquid Sampler into a splitless inlet of a 7890A gas chromatograph. Between samples, the syringe of the automatic liquid sampler was cleaned by flushing 10 times with acetone and 10 times with hexane. The GC was equipped with an Agilent DB-WAXetr column of 30 m x 0.25 mm x 0.25 µm coupled with a flame ionization detector (FID) at 250°C. The program was as follows: 2 minutes on 60°C then temperature was increased by 30°C/minute to 180°C, followed by an 5°C/minute increase of temperature to 230°C. Between samples, the column was heated to 245°C for 15 minutes. To check the retention times and identify the compounds in the extracts, 2–4 µl of a blend of authentic standards of all pheromone compounds (1:1:1:1) was injected before and after every series of injections. Authentic standards used were methyl salicylate (Reagent Plus® ≥99% GC, Sigma Aldrich, US), benzyl cyanide (analytical standard >98%, Sigma Aldrich, US), 2-methoxyphenol (guaiacol) (pure reference standards (1 gram) in solid form, Sigma Aldrich, US), and indole (pure reference standards (1 gram) in solid form, Sigma Aldrich, US). Guaiacol and indole were first dissolved in hexane before injected. Areas under the pheromone peaks were determined using Agilent ChemStation (version B.04.03) .

For every pheromone sample, net amount of each compound was calculated relative to 200 ng pentadecane internal standard. The quantity from all virgin and mated females from Wageningen *P. napi* were used to construct box plots in SPSS v28(Mann Whitney U tests, Table S2) For the heat maps of both species and populations, we calculated the average quantity for each compound using all individuals sampled (Table S3 and S4), and heatmaps were visualized by Biorender. The phylogenies of the *P. napi* populations were constructed using the CO1 barcodes of representatives from each population downloaded from BOLD systems analysis. The accessions used for each population were: Abisko - MW502229; Stockholm - EULEP6041-20; Wageningen- GU669668; Jura- MN141461; bryoniae- MN141756; Costa Brava- GU669673; and Pyerenese - GU669672. Chemical hierarchical clustering was performed in ChemMine tools (2011) using AP Tanimoto values for each compound and hierarchical clustering was performed with default settings.

To create the PCA of the wild-caught *P. napi* from different populations, the quantities of each of the four compounds for each individual were used (Table S4). The Principal Component Analysis (PCA) in R using the `vegan` package (Dixon et al, 2003) was used with the grouping variable was coded as a factor. The PCA was conducted using the `rda()` function with the arguments `center = TRUE` and `scale = TRUE`. Two criteria were used to determine the number of principal components retained for interpretation: Eigenvalues greater than 1 were considered significant and retained. A biplot of the PCA was created using `ordiplot()` and `points()`, displaying sites (observations) with symbols and colors representing group membership. Additionally, we fitted vectors of the odor compounds onto the PCA plot using the `envfit()` function with 10,000 permutations, visualizing only those variables with a significance threshold of $p < 0.001$.

**Table S2** Mated versus virgin *P. napi* Wageningen samples

| Odor | Mated mean +- sd ng | N mated | Virgin mean +- sd ng | N virgin | Test | U | P-value |
|---|---|---|---|---|---|---|---|
| Methyl salicylate | 69.14 +-63.17 | 10 | 19.88 +- 26.97 | 21 | Mann-whitney U | 12 | <0.001 |
| Guaiacol | 11.69 +- 10.49 | 10 | 4.21 +- 6.52 | 21 | Mann-whitney U | 9.5 | <0.001 |
| Benzyl cyanide | 102.15 +- 108.37 | 10 | 25.7 +-43.72 | 21 | Mann-whitney U | 6.5 | <0.001 |
| Indole | 40.57 +- 30.21 | 10 | 23.71 +- 23.42 | 21 | Mann-whitney U | 38 | 0.001 |

**Table S3)** Pieridae post-mating odors samples

| Species | Locations | N | Mean ng +- SD MesA | Mean ng +- SD Guaia | Mean ng +- SD BC | Mean ng +- SD Indole |
|---|---|---|---|---|---|---|
| *Pieris napi* | WAG | 16 | 67.29 +- 72.8 | 9.84 +- 9.44 | 49.34 +- 64.47 | 33.17 +- 48.33 |
| *Pieris bryoniae* | JUR | 13 | 4.21+- 3.17 | 0 | 4.44+- 3.51 | 0 |
| *Pieris rapae* | WAG, COB, PYR | 5 | 3.97 +- 4.25 | 0 | 0.79+-0.83 | 32.9 +- 22.8 |
| *Pieris mannii* | WAG | 8 | 9.89+-12.38 | 0 | 0 | 15.86+-10.96 |
| *Pieris brassicae* | WAG | 4 | 0.82 +- 0.99 | 0.85 +- 1.71 | 36.99 +- 48.7 | 0 |
| *Pieris cheiranth-* | La Palma | 4 | 25.19 +- 9.99 | 3.29 +- 1.9 | 2.9 +- 2.87 | 0 |
| *Pontia daplidice* | PYR | 6 | 2.86 +- 5.81 | 4.62 +- 2.90 | 0.88 +- 1.61 | 0 |
| *Anthocharis cardamine* | WAG | 2 | 25.84 +- 16.45 | 1.65 +- 0.74 | 0.94 + 1.32- | 0 |
| *Euchloe eversi* | Tenerife | 2 | 15.72 +-1.79 | 0 | 0.43 +- 0.24 | 0 |
| *Aporia crataegi* | JUR | 3 | 2.43 +- 1.99 | 0 | 1.81 +-1.411 | 0 |
| *Colias croceus* | COB | 7 | 1.99 +- 2.57 | 4.84 +- 3.48 | 0 | 0 |
| *Colias palaeno* | ABS | 4 | 1.81 +- 2.72 | 3.48 +- 6.30 | 0 | 0 |
| *Leptidea sinapis* | STK, JUR | 11 | 0.65 +- 0.93 | 5.01+- 3.88 | 0 | 0 |

**Table S4***) Pieris napi* populations post-mating odors

| Locations | N odor samples | N mating rate | Mean ng +- SD MesA | Mean ng +- SD Guaia | Mean ng +- SD BC | Mean ng +- SD Indole |
|---|---|---|---|---|---|---|
| ABS | 18 | 25 | 42.01+- 58.29 | 2.63+- 1.4 | 11.68+- 15.39 | 12.77 +- 10.89 |
| STK | 15 | 2 | 25.87 +- 75.35 | 2.75 +- 2.31 | 31.65 +- 57.31 | 20.87 +- 11.12 |
| WAG | 16 | 16 | 67.29 +- 72.8 | 9.84 +- 9.44 | 49.34 +- 64.47 | 33.17 +- 48.33 |
| JUR | 8 | 12 | 2.78 +- 1.07 | 0 | 5.88 +- 8.35 | 6.12 +- 14.2 |
| BRY | 13 | 5 | 4.21 +- 3.17 | 0 | 4.44 +- 3.51 | 0 |
| COB | 14 | 18 | 19.74 +- 49.13 | 2.86 +- 3.3 | 13.02 +- 17.65 | 15.93 +- 8.85 |
| PYR | 10 | 3 | 4.04 +- 2.61 | 6.27 +- 3.15 | 6.23 +- 6.09 | 0 |

**Table S5)** *Pieris napi* and *Trichogramma* sampled per location

| Location | 2022 | | | | 2023 | | | | Phoretic *trichogramma*? | Total #wass / P. napi |
|---|---|---|---|---|---|---|---|---|---|---|
| | # P. napi adults | # eggs | # *Trichogramma* | Sampling days | # P. napi adults | # eggs | # *Trichogramma* | Sampling days | | |
| ABS | 224 | 630 | 7 | 9 | 31 | 85 | 5 | 3 | N | 0.047 |
| STK | 32 | 0 | 1 | 9 | 26 | 46 | 1 | 2 | Y (1) | 0.017 |
| WAG | 62 | 119 | 32 | 11 | 7 | 0 | 13 | 2 | Y (3) | 0.65 |
| JUR | 10 | 10 | 0 | 2 | 21 | 81 | 0 | 5 | N | 0 |
| BRY | 19 | 10 | 0 | 2 | 61 | 81 | 0 | 5 | N | 0 |
| COB | 115 | 355 | 59 | 9 | 38 | 214 | 25 | 5 | Y (2) | |
| PYR | 26 | 14 | 0 | 5 | 26 | 10 | 0 | 2 | N | 0 |

#### S4 Genome editing and mutant screening

##### *S4.1 OR45b identification and phylogeny*

The identification of OR45b as a conserved receptor in *Pieris*, described in detail in Wang *et al.* (2023). The protein sequence of OR45b used in this study was obtained from Wang *et al. (2023)*, who identified olfactory receptor (OR) genes in *Pieris brassicae* through transcriptomic analysis of larval head tissues. Briefly, they performed de novo RNA sequencing of L3 caterpillar heads and quantified OR expression across developmental stages using qPCR. Total RNA was isolated from pooled larval and adult antennae, and transcriptomes were assembled by mapping reads to the *P. brassicae* genome (GCA_942653925) using Hisat2 and StringTie. Candidate chemoreceptors were identified through homology searches (DIAMOND) using known OR sequences from related *Pieris* species, and expression levels were calculated as transcripts per million (TPM). OR45b was among the deorphanized receptors identified in this dataset. The amino acid sequence for OR45b retrieved from this dataset was used as the reference in our BLAST searches, phylogenetic analyses, and subsequent CRISPR knockout experiments.

The phylogenetic tree was constructed in Geneious prime 2023.0.4using the amino acid sequences of OR45b and OR45a from Wang *et al.*, 2023 for *Pieris brasice, P. rapae, P. napi*, and *P. macdonoughii*. *P. brassicae* ORCO was used as the outgroup. The sequence of OR45b from P. brassicae was input into NCBI protein blast filtered for all Pieridae and all Nymphalidae separately. The top 50 matches from both the Pieridae and Nymphalidae results were aligned with the eight Pieris OR45a and b sequences and the ORCO outgroup with MAFFT (Katoh *et al.*, 2009) with default settings, and trimmed to remove any partial sequences and perfect alignment at both ends. A neighbor-joining tree with 1000 bootstraps was used to construct a consensus tree with consensus values at the nodes.

The tree was edited in Figtree to denote the taxonomic classification of the species from which the sequences came, and the clades of olfactory receptors named by the receptors in the clade which had already been de-orphanized in at least one of the species included in the clade.

##### *S4.2 CRISPR*

The first exon of *OR45b* was targeted to knock out the gene. The sgRNA was designed with Geneious Prime v10.0.9 (Geneious, New Zealand) by searching -17bp-NGG sequences. The designated sgRNA was evaluated by Exonerate 2.0 [38] searching against the genome by the est2genome function with a loose cut-off of 50 scores. sgRNA template was synthesized by mixing 5 µl of sgRNA primer (ATTTAGGTGACACTATATTTTTGGTGTTCCATGGCCGTTTTAGAGCTAGAAATAGCAAG), 5 µl of constant primer (AAAAGCACCGACTCGGTGCCACTTTTTCAAGTTGATAACGGACTAGC

CTTATTTTAACTTGCTATTTCTAGCTCTAAAAC), 50 µl of Q5 Hot Start High-Fidelity 2 × Master Mix (NEB, USA) as well as 40 µl of $H_2O$, incubated at 98 °C for 2 min, followed by 35 cycles each at 98 °C for 20 s, 65 °C for 10 s and 72 °C for 10 s; the program ends at 72 °C for 5 min. The cycling product was purified as sgRNA template using QIAquick PCR Purification Kit (Qiagen, Germany) according to the manufacturer's instructions. sgRNA was synthesized using a MEGAscript™ SP6 Transcription Kit (Invitrogen, USA) and incubated at 37 °C for four hours followed by adding 1 µl Turbo DNase and incubating at 37 °C for 15 min. The reaction products were purified with Monarch RNA Cleanup Kit. The concentration of sgRNA was determined by DeNovix (DeNovix, USA).

Before injection, a mixture of 2 µl of 300 ng/µl sgRNA, 1 µl of EnGen Spy Cas9 NLS (NEB, USA), 1 µl of 10 × NEBuffer r3.1 and 6 µl of $H_2O$ was incubated at 25 °C for 10 min. sgRNA/Cas9 mixture was colored with 1 µl food dye as a visual marker for injected eggs before loading the mixture into the glass needle of FemtoJet (Eppendorf, Germany). Newly laid 30-min-old eggs were collected from our laboratory colony and were injected with colored sgRNA/Cas9 complex. The injected eggs were incubated at 25 °C for four-five days until hatching. Around 100 caterpillars were reared on *B. oleracea* until pupation.

##### *S4.3 Screening*

Following eclosion of the butterflies, one leg of each butterfly was then dissected to screen for mutants. The leg samples were lysed and extracted using MyTaq Extract-PCR Kit (Bioline, UK) according to manufacturer's instructions. Two microliters of extract as gDNA template were added into the PCR reaction system of 12.5 µl of 2 × MyTaq Red Mix (Bioline, UK), 2 µl of forward primer (ATGGCTAACATATCTGATACTTTTAATATC) and reverse primer (TTACTGCAATACCTTCTCATAATGCC) each and 6.5 µl of $H_2O$. The reaction system was incubated at 95 °C for 2 min followed by 38 cycles each at 95 °C for 15 s, 55 °C for 15 s and 72 °C for 10 s; the program ends at 72 °C for 2 min. The PCR products were checked by gel electrophoresis and sequencing (Eurofins, The Netherlands). The transformed butterflies are further referred to as "knockout" or KO and the butterflies from the same rearing that did not undergo genome editing are referred to as "wildtype" or WT. The resulting mutation be found in Figure S3.

#### S5 Electrophysiology

##### *S5.1 Dose-response analysis*

To assess olfactory sensitivity across species and sexes, we performed electroantennogram (EAG) recordings on adult males and females of *Pieris brassicae*, *P. mannii*, *P. rapae*, and *P. napi*. The excised antenna was mounted between two glass capillaries filled with EAG Ringer solution (as per standard insect saline recipes), with each capillary containing a silver/silver chloride (Ag/AgCl) electrode. One electrode was connected to ground, and the other to a 10× preamplifier (Ockenfels Syntech, Germany). Signals were digitized using an IDAC-2 acquisition controller (Ockenfels Syntech) and recorded with EAG Pro software. The antenna was oriented ventral side up to maximize exposure to odor pulses.

Odor stimuli consisted of five compounds: methyl salicylate, benzyl cyanide, indole, cis-3-hexenyl acetate, and linalool (≥98% purity Sigma Aldrich). Odorants were diluted in HPLC-grade hexane to create a three-step dose series: 0.01, 0.1, and 1.0 (v/v). A 10 µL aliquot of each solution was pipetted onto a strip of filter paper inserted into a disposable pipette tip, which was then loaded into the stimulus cartridge. Each odorant was tested at all three concentrations per individual. Stimuli were presented as 0.5–1.0 s air pulses delivered via a continuous humidified air stream (1 L/min flow rate). The first puff from each newly loaded stimulus cartridge was discarded to avoid solvent front effects, and responses were recorded beginning with the second puff.

Odorant presentation order was randomized across individuals to reduce position and fatigue effects; however, concentrations were always presented in ascending order. At least 20 seconds elapsed between stimulus puffs to allow antennal recovery and baseline stabilization. Each trial included multiple control stimuli: clean air, blank filter paper, filter paper with only hexane, and empty pipette tips. All EAG responses were recorded in mV and baseline-corrected and normalized relative to the individual's response to hexane alone. The effect of dose on relative response for each species, sex, and odor was tested for significance using a linear regression with concentration as a predictor in SPSS version 28.

##### *S5.2 Electroantennogram (EAG) recording of mutants versus wildtype*

A panel of 47 chemical compounds that are potentially involved in the tritrophic interaction including esters, alcohols, aldehydes, alkene, ITCs, heterocyclics and nitriles (Table S1) were selected as candidate ligands (concentration at $10^{-2}$ v/v) of OR45b based on previous literature. EAG responses were investigated for the left antennae of three-day-old males and females (n = 17-20 for both genotypes). Filter papers were inserted into Pasteur pipettes and loaded with 10 µl of a certain chemical diluted in paraffin oil, uncontaminated filter paper and paraffin oil-loaded filter paper was employed as negative controls. The left antennae of butterflies were excised at the basal end and the distal tip was also removed for better conductivity. Both antenna ends were placed in glass capillaries filled with EAG ringer solution. Ag/Ag-Cl wires were inserted into both capillaries, with one connected to a ground electrode and one connected to a 10× pre-amplifier (Ockenfels Syntech, Germany), to form an electric circuit. The electrical signals were converted from analog to digital using an IDAC-2 (Ockenfels Syntech, Germany) and sent to a personal computer. The first puff of all newly loaded chemicals was discarded and responses were recorded starting with the second puff of chemicals, the magnitude of response was recorded in mV. At least 20 seconds passed between odor puffs, to allow the antenna to recover and the signal to return to a stable baseline. Compounds were tested in random orders. The response signals were recorded using EAG Pro software (Ockenfels Syntech, Germany). The response of each chemical was calibrated by subtracting the background signal (response to uncontaminated filter paper). Differences of EAG responses to the chemicals between the two genotypes were tested by using the Student's t-test when the values were normally distributed and had similar variance, or tested by using the Kruskal-Wallis test when the data failed to meet the criteria of the Student's t-test. The results from the full panel can be found in Figure S3.

**Table S1.** Chemical compounds that were used for the electroantennographically test

| Name | CAS number | Purity | Manufacturer |
|---|---|---|---|
| Benzoic acid | 65-85-0 | ≥99.5% | Sigma-Aldrich |
| (*E*)-Anethole | 4180-23-8 | 99.0% | Sigma-Aldrich |
| 1-Pentanol | 71-41-0 | ≥ 99.0% | Sigma-Aldrich |
| (*Z*)-2-Penten-1-ol | 1576-95-0 | 95.0% | Sigma-Aldrich |
| 1-Hexanol | 111-27-3 | 98.0% | Fluka |
| (*Z*)-3-Hexen-1-ol | 928-96-1 | 98.0% | Sigma-Aldrich |
| 1-Octen-3-ol | 3391-86-4 | 98.0% | Sigma-Aldrich |
| Geraniol | 106-24-1 | 98.0% | Sigma-Aldrich |
| (*E*)-2-Hexen-1-ol | 928-95-0 | ≥ 95.0% | Sigma-Aldrich |
| Linalool | 78-70-6 | 97.0% | Sigma-Aldrich |
| 1,8-Cineole | 470-82-6 | 99.0% | Sigma-Aldrich |
| 1-Penten-3-ol | 616-25-1 | 99.0% | Sigma-Aldrich |

**Table S1.** *Continued*

| Name | CAS number | Purity | Manufacturer |
|---|---|---|---|
| Phenylethyl alcohol | 60-12-8 | ≥99.0 % | Fluka |
| 1-Methoxy-2-propanol | 107-98-2 | ≥ 99.5% | Sigma-Aldrich |
| 3-Pentanol | 584-02-1 | 98.0% | Sigma-Aldrich |
| 3-Methyl-1-butanol | 123-51-3 | ≥ 99.0% | Sigma-Aldrich |
| 3-Octanol | 589-98-0 | 99.0% | Sigma-Aldrich |
| Hexanal | 66-25-1 | 98.0% | Sigma-Aldrich |
| (*E*)-2-Hexenal | 6728-26-3 | 98.0% | Sigma-Aldrich |
| Benzaldehyde | 100-52-7 | ≥99.0% | Fluka |
| Phenylacetaldehyde | 122-78-1 | ≥95.0% | Fluka |
| Heptaldehyde | 111-71-7 | 95.0% | Sigma-Aldrich |
| 1-Nonanal | 124-19-6 | 95.0% | Sigma-Aldrich |
| 3-Methylbutanal | 123-51-3 | ≥ 98.5% | Fluka |
| 2-Methylbutanal | 96-17-3 | 95.0% | Sigma-Aldrich |
| (*Z*)-2-Pentenal | 1576-87-0 | 95.0% | Sigma-Aldrich |
| Nonane | 111-84-2 | 99.0% | Sigma-Aldrich |
| Limonene | 5989-27-5 | 97.0% | Sigma-Aldrich |
| *α*-Pinene | 80-56-8 | 98.0% | Sigma-Aldrich |
| *β*-Pinene | 127-91-3 | ≥ 95.0% | Sigma-Aldrich |
| Myrcene | 123-35-3 | ≥ 99.0 | Sigma-Aldrich |
| *β*-Caryophyllene | 87-44-5 | ≥ 98.0% | Sigma-Aldrich |
| Pentyl acetate | 628-63-7 | 99.0% | Sigma-Aldrich |
| Hexyl acetate | 142-92-7 | 99.0% | Sigma-Aldrich |
| (*Z*)-3-Hexenyl acetate | 3681-71-8 | 98.0% | Sigma-Aldrich |
| Methyl salicylate | 119-36-8 | ≥ 98.0% | Sigma-Aldrich |
| Indole | 120-72-9 | ≥ 99.0% | Sigma-Aldrich |
| Benzothiazole | 95-16-9 | ≥ 96.0% | Sigma-Aldrich |
| Allyl ITC | 57-06-7 | ≥ 95.0% | Sigma-Aldrich |
| Benzyl ITC | 622-78-6 | 98.0% | Sigma-Aldrich |
| Methyl ITC | 556-61-6 | 97.0% | Fluka |
| Butyl ITC | 592-82-5 | 99.0% | Sigma-Aldrich |
| Phenyl ITC | 103-72-0 | 98.0% | Sigma-Aldrich |
| 3-Hydroxy-2-butanone | 513-86-0 | ≥ 98.0% | Sigma-Aldrich |
| 3-Octanone | 106-68-3 | ≥ 98.0% | Sigma-Aldrich |
| Benzyl cyanide | 140-29-4 | 98.0% | Sigma-Aldrich |
| Paraffin oil | 8012-95-1 | - | Sigma-Aldrich |

#### S6 Behavior assays

##### *S6.1 Y-tube*

To assess the attraction of *Trichogramma* to methyl salicylate, bioassays with three strains of *T. a evanescens*, *T. semblidis*, and *T. cacoeciae*, all collected in Wageningen, were conducted using a Y-tube olfactometer. similar to the one described by Fatouros et al.(2014). Air was filtered through activated charcoal and humidified by passing through a bottle with tap water before entering the system. The air was split into two, each tube leading into a glass container (500 ml) with an odour source of either a virgin or mated female from Wageningen as described in section 1.3.2. From the glass containers containing the odour source a tube led into one of the arms of a glass Y-tube olfactometer (stem cm, arms cm, cm). The Y-tube was mounted on a board with an incline so that the wasps would walk upwards. The airflow was set at 300 mL min−1 in each arm using flow meters. The bioassays were carried out in the laboratory of Wageningen University between 11:00 and 17:00 at room temperature. One diffused uniform light source was situated above the Y-tube set-up and was kept centred directly over the y-tube.

The butterflies were introduced in the glass container which were shut using a Viton O-ring and a metal clamp. After introducing the butterflies in the containers, their odor was allowed to distribute through the system for 2 minutes. Air flow first passed through an activated carbon filter, then through distilled water and finally entered the Y-tube at 300 ml/min. The olfactometer was thoroughly washed with ethanol, rinsed with water, and air-dried at room temperature after every test day.

Naïve female wasps of 3-5 days old were well-fed and had not yet had oviposition experience. 1 hour before testing, female wasps were separated from the rearing and kept in plastic tubes with a drop of honey. Ten female wasps were released in the glass Y-tube olfactometer (Table S6.6). They could select to walk toward the side of the Y-tube with air flow from the glass container housing two mated female butterflies vs. the side of the Y-tube with two virgin females. After 20- 30 minutes, the wasps collected in each of the trapping bulbs were counted. When a wasp did not make a choice within 30 minutes, not crossing the redline indicted in figure S6.1, it was recorded as a “no response” and excluded from the statistical analysis. Each wasp was used only once to test for innate response. To exclude any bias and directional effects, the position of the tubes leading from the glass containers containing the odour source to the Y-tube olfactometer, was exchanged after every 2nd trial.

In the following assays (Table S6.7), these methods were repeated using *Trichogramma cacoeciae* from Wageningen were tested between two odor sources: clean air or mated butterflies from one of the populations, either WAG, COB, JUR, or PYR. They were tested using the same protocol described above.

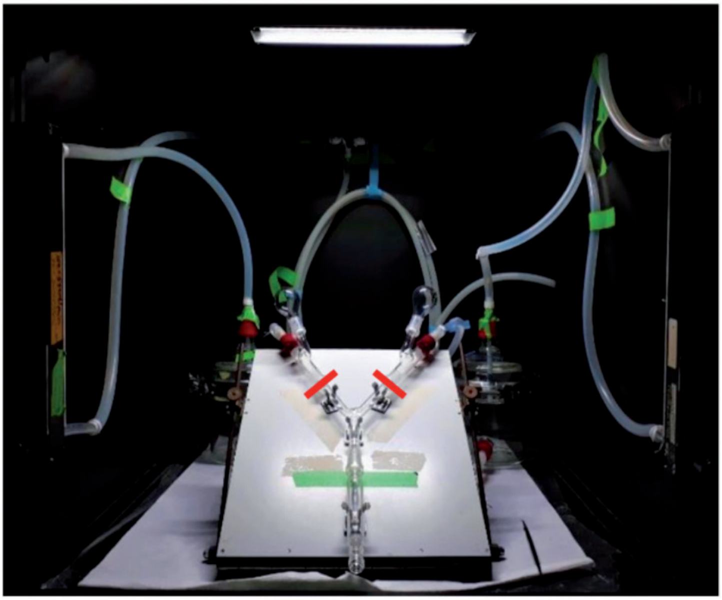

**Figure S2)** Y-tube olfactometer set-up. Test odors collect in bulbs, towards which wasps may walk. Red bars represent the divide between "response" and "no response".

##### *S6.2 Static olfactometer*

In order to explore the influence of *OR45b* on mating frequency and consequently on attractiveness of both butterfly genotypes to *T. evanescens* w a sps, the following experiment was conducted: we used either freely mated or virgin WT butterfly females and freely mated or virgin KO butterfly females as odor sources in this experiment. Similar to the oviposition experiment (1), butterflies were allowed to mate freely with others of the same genotype. All butterflies were placed in mesh cages immediately after eclosion. For the mated butterfly treatment, 15 female butterflies were placed in a single cage together with 30 males of the same age and genotype and allowed to mate freely without cabbage plant. For the virgin female treatment, 15 newly eclosed virgin females were placed in a cage together and not allowed to have contact with males for 10 days before testing. Tests were performed 10 days after eclosion with both mated and virgin WT or KO females. Mating rates for butterflies in these tests can be found in Figure S6.9.

To investigate behavioral responses of the wasps to butterfly odors, we used a two-chamber olfactometer (18 cm high × 12 cm in diameter) described previously (Faoutors et al., 2005, Huigens et al., 2009). Two butterflies from one of the four treatments were placed in one chamber of the olfactometer while the other chamber was kept empty. We specifically used two butterflies; they would fly and trigger each other to exhibit the "mate-refusal posture", potentially emitting benzyl cyanide (Andersson et al., 2003). Data was only used when mated butterflies adopted the mate-refusal posture at least once per wasp trial. Butterflies were placed at the bottom of the chamber and the top was covered with fine mesh. On top of the mesh we placed a single, mated and honey-fed female *T. evanescens* that had eclosed within the previous 3 days and had no prior oviposition experience. To close the olfactometer, a glass lid was placed on top so that the parasitoid was able to walk around on the mesh and on the lid above the two chambers. The wasp was released in the center between the two chambers, and a 300 second timer was started immediately. The number of seconds that the parasitoid spent on the side of the chamber with butterflies within 300 seconds was recorded. Each but-terfly pair (n = 9 for both WT and *OR45b* KO mated butterflies, n = 10 for WT virgin and n = 12 for *OR45b* KO virgin butterflies) was tested with three individual egg parasitoids, with each wasp (n = 27-30) being used for only a single trial. Between each of the three trials, the olfactometer was rotated 90 degrees. After three trials, a new butterfly pair was used from a randomly selected treatment. The chamber was opened to allow the air within to be refreshed between butterfly pairs. T he difference of egg parasitoid preference to the butterflies among the treatments was tested using one-way ANOVA followed by Tukey's post-hoc test.

***S6.4 Mating rates in the field***

From each population, 5 – 10 butterflies used for odor collection were frozen and later dissected using a scalpel and pointy forceps. Wings and head were discarded. The abdomen was carefully opened and the bursa copulatrix was isolated. With pointy forceps, the bursa copulatrix was carefully opened, and spermatophores were isolated. Mating frequency was established by counting the number of spermatophores present in the bursa copulatrix.

***S6.5 Egg laying dynamics of mutant versus wildtype butterflies***

To compare the egg-laying behavior of the two genotypes, we designed three different experiments. The results are in Figure S6.

(1) In order to determine whether *OR45b* has an effect on mating and egg-laying behavior, a single pair of newly emerged male and female butterflies of the same genotype were put into individual cages (40 cm × 40 cm ×60 cm) so that the butterflies could mate freely during the experiment (n = 14 for both genotypes). The pairs were supplied with a four-week-old Brussels sprouts plant for oviposition and honey water for food. The butterflies were allowed to mate freely and repeatedly and were exposed to the same plant during the entire experimental period of eight days. The number of eggs per butterfly pair was counted daily for seven days from Day 0 to Day 7, with the day that butterflies

were placed in the cages marked as Day 0. The eggs were allowed to hatch over the following week in order to measure the hatching rate and to determine whether or not the eggs had been fertilized. Immediately at the end of the eighth day, the butterflies were collected and frozen for dissection. The spermatophores were removed and counted from each butterfly to compare the number of times both WT females and KO females mated during eight days. All female butterflies were collected and frozen at -20 °C. The frozen butterflies were then dissected under 20× magnification to count the number of spermatophores present in the bursa copulatrix, as the number of spermatophores is equivalent to the number of successful copulations. The bursa copulatrix was removed from the abdomen and carefully opened so that the spermatophores within it could be counted (Figure S8). The fullness of the spermatophore was ranked as follows: a. full, b. partially full, or c. empty. A spermatophore was considered as full (a) when its envelope was taut due to the high volume of its contents which completely filled the envelope. This state indicates that copulation was recent, as spermatophore digestion begins approximately 24 hours post mating. An empty spermatophore (c), for which only the envelope is present and the contents within have been fully digested and thus are no longer visible, indicates that copulation had occurred at least two days prior to dissec-tion. Anything in between these states was considered partially full (b).

(2) In order to determine if the effect of the knockout on oviposition was linked to differences found in mating frequency, we compared the number of eggs laid by couples of the same genotype when butterflies were only allowed to mate once. To control for mating frequency, three-day-old virgin females were put together with three-day-old virgin males in a cage (40 cm × 40 cm ×60 cm). Upon mating, pairs were moved to a new cage, and once they separated, the male was removed entirely. Therefore, each cage contained either a single WT female (n = 17) or KO female (n = 19) that had mated once. Then each mated female butterfly was placed in a cage supplied with a cabbage plant and honey-water as described above. The number of eggs laid per female and the number of newly hatched caterpillars were counted every day.

(3) Finally, in order to determine whether effects on egg laying and mating resulted from changes in the KO female, or the KO male, or both, we further crossed the mutant butterflies and wildtype butterflies and conducted the same experiment as above. Each cage (40 cm × 40 cm ×60 cm) had a single pair that consisted of either a WT female and a KO male (n = 10) or a KO female and a WT male (n = 10), and butterflies were allowed to mate freely during the experiment. The female butterflies were collected and dissected, and the number of eggs per pair and newly hatched caterpillars was counted as we described above.

In all three experiments, differences in egg-laying dynamics between genotypes (WT and KO) or treatments (cross-mating) were tested by using a general linear model (GLM) with negative binominal distribution. Differences of number of eggs hatching, egg hatching rate and number of matings between genotypes (WT and KO) or treatments (cross-mating) were tested by using Student's t-test when the date was normally distributed and has equal variance, otherwise differences were tested using the Wilcoxon rank-sum test when the criteria of the Student's t-test did not apply.

##### *S6.6 Oviposition choice of mutant versus wildtype butterflies*

An oviposition preference assay was conducted to evaluate whether WT and KO female butterflies respond differently to the odors of egg-infested host plants (Figure S7). In short, *Brassica oleracea* var. *gemmifera* Cyrus plants were infested with 7 - 10 egg clutches by WT butterflies. Singly mated butterflies of both genotypes were prepared 24-48 hours prior to the behavioral test as described above for oviposition experiment (2). Plants were infested with eggs 72 hours beforehand to obtain infested plants before starting the tests. Just prior to the test, the egg clutches were gently removed from the plants (subsequently called egg-infested) and any possible damage caused by this removal was mimicked on the control plant (called non-infested). The two plants were grown from the same batch of seeds and under the same environmental conditions. A previously egg-infested and a non-infested plant were then placed at the far end of a mesh cage (1.8 m ×1.8 m ×1.8 m). A female butterfly was released from the opposite side of the cage halfway between the two plants. The number of seconds until the butterfly landed on a plant and the plant identity, were recorded. Then, the number of seconds until an egg was laid on a certain plant, was also recorded. If no egg was laid within 15 minutes, the butterfly was considered a non-responder. Butterflies (n = 41 of each genotype) were tested one time on a pair of plants (n = 18), and each pair of plants was used for a maximum of 6 butterflies. The side of the cage which had the treatment or control plant was alternated between every butterfly tested, the genotype of butterflies was also alternated between WT and KO females. The difference in the time that butterflies spent on selecting a cabbage plant for oviposition was tested by Wilcoxon rank-sum test. The difference in butterfly oviposition-site selection between the two genotypes was tested by using a Chi-square test.

##### *S6.7 Predation*

The experiments were conducted in the same cages in which the great tits (*Parus major*) were housed (Figure S3). Prior to the start of each trial, external feeder trays and water bathing dishes were removed. A wooden cover plate was inserted and placed on top of the cage bedding to reduce potential external influencing factors (e.g., residual food in the bedding) and to ensure that only the test objects were present in the environment.

To assess whether great tits exhibited a preference for butterflies lacking the odor benzyl cyanide, a two-choice assay was employed. In each trial, birds were presented with one control butterfly (treated with hexane only) and one treated butterfly (hexane plus benzyl cyanide at one of two concentrations). The butterflies used were recently frozen wild-type virgin females from the rearing and treated according to the following protocols: 1. Control: A virgin female pipetted with 10 µL of hexane. 2. Treatment: A virgin female pipetted with 10 µL of a benzyl cyanide solution in hexane. The total amount of benzyl cyanide applied ranged between 0.01 µg and 0.1 µg. All solutions were applied to the thorax and abdomen.

Butterflies were placed next to each other in the cage, equidistant from the back and front walls. Each was positioned in a small petri dish containing non-toxic green polymer oven-bake clay (DécoTime) as a background for contrast and to record peck marks from the birds. The hindwings were gently pressed into the clay to hold the butterfly in an upright position, displaying both fore- and hindwing spots in a typical Pieridae deterrence posture. Upon introduction of the petri dishes into the cage, researchers exited the room to minimize disturbance. Bird behavior was recorded using a GoPro Hero 5 (GoPro Inc., San Mateo, USA) camera at 25 fps and 720p resolution. Each bird was given five minutes to interact with the butterflies. Following the trial, any butterfly remains were removed and stored at −20 °C for later analysis.

Each bird underwent two trials across the course of the experiment, conducted on alternating days using the same setup and treatment combinations. To control for side bias, the positions of the treatment and control butterflies were switched between trials. The bird's first choice was recorded as the first butterfly taken into the beak. Birds that did not make a choice within the trial period were excluded from further analysis. The latency to attack (time until the butterfly was taken into the beak) was recorded as a proxy for repellence.

All procedures were conducted in accordance with ethical standards. No harm was inflicted on the great tits, and no invasive or unethical interventions were performed. The license for the study was issued under number NIOO 23.07 (AVD 80100 2019 9005) / IvD 3050 on 07-11-2023.

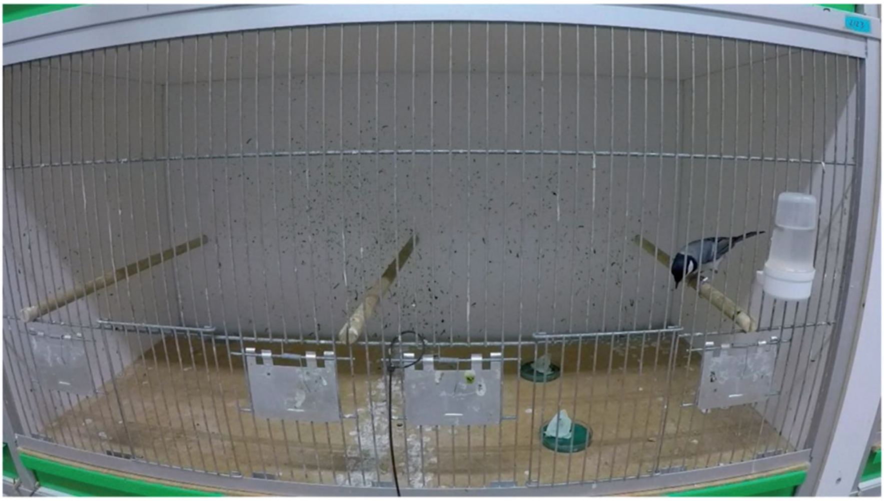

**Figure S3)** Experimental set up for predator choice assays with *Parus major* and dead, virgin *Pieris brassicae* with or without benzyl cyanide applied to abdomen.

### Supplementary Results

#### S7 Post-mating odor emission is not species-specific in Pieridae (Figure 1)

**Table S2** Mated versus virgin *P. napi* Wageningen samples

| Odor | Mated mean +- sd ng | N mated | Virgin mean +- sd ng | N virgin | Test | U | P-value |
|---|---|---|---|---|---|---|---|
| Methyl salicylate | 69.14 +-63.17 | 10 | 19.88 +- 26.97 | 21 | Mann-whitney U | 12 | <0.001 |
| Guaiacol | 11.69 +- 10.49 | 10 | 4.21 +- 6.52 | 21 | Mann-whitney U | 9.5 | <0.001 |
| Benzyl cyanide | 102.15 +- 108.37 | 10 | 25.7 +-43.72 | 21 | Mann-whitney U | 6.5 | <0.001 |
| Indole | 40.57 +- 30.21 | 10 | 23.71 +- 23.42 | 21 | Mann-whitney U | 38 | 0.001 |

**Table S3)** Pieridae post-mating odors samples

| Species | Locations | N | Mean ng +- SD MesA | Mean ng +- SD Guaia | Mean ng +- SD BC | Mean ng +- SD Indole |
|---|---|---|---|---|---|---|
| *Pieris napi* | WAG | 16 | 67.29 +- 72.8 | 9.84 +- 9.44 | 49.34 +- 64.47 | 33.17 +- 48.33 |
| *Pieris bryoniae* | JUR | 13 | 4.21+- 3.17 | 0 | 4.44+- 3.51 | 0 |
| *Pieris rapae* | WAG, COB, PYR | 5 | 3.97 +- 4.25 | 0 | 0.79+-0.83 | 32.9 +- 22.8 |
| *Pieris mannii* | WAG | 8 | 9.89+-12.38 | 0 | 0 | 15.86+-10.96 |
| *Pieris brassicae* | WAG | 4 | 0.82 +- 0.99 | 0.85 +- 1.71 | 36.99 +- 48.7 | 0 |
| *Pieris cheiranth-* | La Palma | 4 | 25.19 +- 9.99 | 3.29 +- 1.9 | 2.9 +- 2.87 | 0 |
| *Pontia daplidice* | PYR | 6 | 2.86 +- 5.81 | 4.62 +- 2.90 | 0.88 +- 1.61 | 0 |
| *Anthocharis cardamine* | WAG | 2 | 25.84 +- 16.45 | 1.65 +- 0.74 | 0.94 + 1.32- | 0 |
| *Euchloe eversi* | Tenerife | 2 | 15.72 +-1.79 | 0 | 0.43 +- 0.24 | 0 |
| *Aporia crataegi* | JUR | 3 | 2.43 +- 1.99 | 0 | 1.81 +-1.411 | 0 |
| *Colias croceus* | COB | 7 | 1.99 +- 2.57 | 4.84 +- 3.48 | 0 | 0 |
| *Colias palaeno* | ABS | 4 | 1.81 +- 2.72 | 3.48 +- 6.30 | 0 | 0 |
| *Leptidea sinapis* | STK, JUR | 11 | 0.65 +- 0.93 | 5.01+- 3.88 | 0 | 0 |

#### S8 Intraspecific variation in *Pieris napi* post-mating odors is shaped by parasitoid pressure rather than geography or genetic distance (Figure 2)

**Table S4)** *Pieris napi* populations post-mating odors

| Locations | N odor samples | N mating rate | Mean ng +- SD MesA | Mean ng +- SD Guaia | Mean ng +- SD BC | Mean ng +- SD Indole |
|---|---|---|---|---|---|---|
| ABS | 18 | 25 | 42.01+- 58.29 | 2.63+- 1.4 | 11.68+- 15.39 | 12.77 +- 10.89 |
| STK | 15 | 2 | 25.87 +- 75.35 | 2.75 +- 2.31 | 31.65 +- 57.31 | 20.87 +- 11.12 |
| WAG | 16 | 16 | 67.29 +- 72.8 | 9.84 +- 9.44 | 49.34 +- 64.47 | 33.17 +- 48.33 |
| JUR | 8 | 12 | 2.78 +- 1.07 | 0 | 5.88 +- 8.35 | 6.12 +- 14.2 |
| BRY | 13 | 5 | 4.21 +- 3.17 | 0 | 4.44 +- 3.51 | 0 |
| COB | 14 | 18 | 19.74 +- 49.13 | 2.86 +- 3.3 | 13.02 +- 17.65 | 15.93 +- 8.85 |
| PYR | 10 | 3 | 4.04 +- 2.61 | 6.27 +- 3.15 | 6.23 +- 6.09 | 0 |

**Table S5)** *Pieris napi* and *Trichogramma* sampled per location

| Loca-tion | 2022 | | | | 2023 | | | | Phoretic *Tricho-gramma*? | Total #wass / P. napi |
|---|---|---|---|---|---|---|---|---|---|---|
| | # P. napi adults | # eggs | # *Tricho-gramma* | Sampling days | # P. napi adults | # eggs | # *Tricho-gramma* | Sampling days | | |
| ABS | 224 | 630 | 7 | 9 | 31 | 85 | 5 | 3 | N | 0.047 |
| STK | 32 | 0 | 1 | 9 | 26 | 46 | 1 | 2 | Y (1) | 0.017 |
| WAG | 62 | 119 | 32 | 11 | 7 | 0 | 13 | 2 | Y (3) | 0.65 |
| JUR | 10 | 10 | 0 | 2 | 21 | 81 | 0 | 5 | N | 0 |
| BRY | 19 | 10 | 0 | 2 | 61 | 81 | 0 | 5 | N | 0 |
| COB | 115 | 355 | 59 | 9 | 38 | 214 | 25 | 5 | Y (2) | |
| PYR | 26 | 14 | 0 | 5 | 26 | 10 | 0 | 2 | N | 0 |

**Table S6)** *Trichogramma* species comparison in y-tube

| Line | Species | N (10 wasps per trial) | Mean % to mated female | One-sample t test | One sided P-value |
|---|---|---|---|---|---|
| 16_ben_20 | *T. evanescens* | 6 | 55.11 | .817 | 0.226 |
| 64_ eng_20 | *T. evanescens* | 5 | 58.15 | .762 | 0.240 |
| 69_eng_20 | *T. evanescens* | 6 | 55.07 | .725 | 0.248 |
| 18_eng_21 | T. semblidis | 6 | 55.73 | .734 | 0.248 |
| 6_gre_21 | T. cacoeciae | 6 | 66.35 | 2.255 | 0.037 |

**Table S7)** *Trichogramma cacoeciae* response to different *Piers napi* populations

| Butterfly popula-tion | N (10 wasps per trial) | Mean % to mated female | One-sample t test | One sided P-value |
|---|---|---|---|---|
| WAG | 16 | 62 | 1.76 | 0.04 |
| JUR | 7 | 50 | 0.04 | 0.97 |
| COB | 10 | 73 | 4.46 | 0.002 |
| PYR | 25 | 50 | .063 | 0.475 |

#### S9 Interspecific variation and genetic basis of olfactory responses to post-mating odors (Figure 3)

**Table S8** Dose response EAG stats linear regression

| odor | species | sex | N (for three concentrations) |
|---|---|---|---|
| BC | *P. brassicae* | male | 27 |
| | *P. brassicae* | female | 39 |
| | *P. napi* | male | 30 |
| | *P. napi* | female | 36 |
| | *P. rapae* | male | 36 |
| | *P. rapae* | female | 27 |
| | *P. mannii* | male | 18 |
| | *P. mannii* | female | 23 |
| MeSa | *P. brassicae* | male | 27 |
| | *P. brassicae* | female | 39 |
| | *P. napi* | male | 30 |
| | *P. napi* | female | 36 |
| | *P. rapae* | male | 35 |
| | *P. rapae* | female | 27 |
| | *P. mannii* | male | 18 |
| | *P. mannii* | female | 24 |
| Indole | *P. brassicae* | male | 27 |
| | *P. brassicae* | female | 38 |
| | *P. napi* | male | 30 |
| | *P. napi* | female | 36 |
| | *P. rapae* | male | 36 |
| | *P. rapae* | female | 27 |
| | *P. mannii* | male | 18 |
| | *P. mannii* | female | 25 |
| Linalool | *P. brassicae* | male | 27 |
| | *P. brassicae* | female | 39 |
| | *P. napi* | male | 30 |
| | *P. napi* | female | 36 |
| | *P. rapae* | male | 34 |
| | *P. rapae* | female | 27 |
| | *P. mannii* | male | 18 |
| | *P. mannii* | female | 24 |

**Table S8** *Continued*

| odor | species | sex | N (for three concentrations) |
|---|---|---|---|
| Cis-3 | *P. brassicae* | male | 27 |
| | *P. brassicae* | female | 39 |
| | *P. napi* | male | 30 |
| | *P. napi* | female | 36 |
| | *P. rapae* | male | 35 |
| | *P. rapae* | female | 27 |
| | *P. mannii* | male | 18 |
| | *P. mannii* | female | 24 |

**Table S9)** Coeffieicents table of regression of dose-response EAGS in *Pieris*

| | | | | | B | Std. Error | Beta | | |
|---|---|---|---|---|---|---|---|---|---|
| bc | female | *P. brassicae* | 1 | (Constant) | 0.287 | 0.115 | | 2.484 | 0.018 |
| | | | | Concentration | 0.900 | 0.199 | 0.597 | 4.527 | **0.000** |
| | | *P. mannii* | 1 | (Constant) | 0.709 | 0.294 | | 2.408 | 0.025 |
| | | | | Concentration | 0.692 | 0.496 | 0.291 | 1.394 | 0.178 |
| | | *P. napi* | 1 | (Constant) | 0.384 | 0.214 | | 1.792 | 0.082 |
| | | | | Concentration | 1.117 | 0.369 | 0.461 | 3.026 | **0.005** |
| | | *P. rapae* | 1 | (Constant) | 0.116 | 0.132 | | 0.881 | 0.387 |
| | | | | Concentration | 1.055 | 0.227 | 0.681 | 4.645 | **0.000** |
| | male | *P. brassicae* | 1 | (Constant) | 0.523 | 0.221 | | 2.368 | 0.026 |
| | | | | Concentration | 1.149 | 0.381 | 0.516 | 3.016 | **0.006** |
| | | *P. mannii* | 1 | (Constant) | 0.375 | 0.199 | | 1.883 | 0.078 |
| | | | | Concentration | 0.810 | 0.344 | 0.508 | 2.358 | **0.031** |
| | | *P. napi* | 1 | (Constant) | 0.490 | 0.164 | | 2.993 | 0.006 |
| | | | | Concentration | 0.722 | 0.282 | 0.436 | 2.560 | **0.016** |
| | | *P. rapae* | 1 | (Constant) | 0.169 | 0.083 | | 2.047 | 0.048 |
| | | | | Concentration | 0.984 | 0.156 | 0.735 | 6.316 | **0.000** |

**Table S9)** *Continued*

| | | | | | B | Std. Error | Beta | | |
|---|---|---|---|---|---|---|---|---|---|
| ci | female | *P. brassicae* | 1 | (Constant) | 0.029 | 0.053 | | 0.548 | 0.587 |
| | | | | Concentration | 0.102 | 0.091 | 0.180 | 1.116 | 0.272 |
| | | *P. mannii* | 1 | (Constant) | 0.313 | 0.246 | | 1.276 | 0.215 |
| | | | | Concentration | 0.771 | 0.423 | 0.362 | 1.821 | 0.082 |
| | | *P. napi* | 1 | (Constant) | 0.197 | 0.111 | | 1.783 | 0.084 |
| | | | | Concentration | 0.267 | 0.191 | 0.233 | 1.399 | 0.171 |
| | | *P. rapae* | 1 | (Constant) | 0.054 | 0.069 | | 0.774 | 0.446 |
| | | | | Concentration | 0.144 | 0.120 | 0.234 | 1.204 | 0.240 |
| | male | *P. brassicae* | 1 | (Constant) | 0.200 | 0.119 | | 1.682 | 0.105 |
| | | | | Concentration | 0.243 | 0.205 | 0.231 | 1.186 | 0.247 |
| | | *P. mannii* | 1 | (Constant) | 0.252 | 0.198 | | 1.273 | 0.221 |
| | | | | Concentration | 0.164 | 0.341 | 0.119 | 0.481 | 0.637 |
| | | *P. napi* | 1 | (Constant) | 0.360 | 0.143 | | 2.521 | 0.018 |
| | | | | Concentration | 0.280 | 0.246 | 0.210 | 1.135 | 0.266 |
| | | *P. rapae* | 1 | (Constant) | 0.222 | 0.058 | | 3.842 | 0.001 |
| | | | | Concentration | 0.157 | 0.107 | 0.247 | 1.465 | 0.152 |
| in | female | *P. bras* | 1 | (Constant) | 0.096 | 0.070 | | 1.359 | 0.182 |
| | | | | Concentration | 0.276 | 0.125 | 0.347 | 2.217 | **0.033** |
| | | *P. mann* | 1 | (Constant) | 0.290 | 0.274 | | 1.056 | 0.302 |
| | | | | Concentration | 0.584 | 0.482 | 0.245 | 1.211 | 0.238 |
| | | *P. napi* | 1 | (Constant) | 0.259 | 0.102 | | 2.530 | 0.016 |
| | | | | Concentration | 0.165 | 0.176 | 0.159 | 0.938 | 0.355 |
| | | *P. rapa* | 1 | (Constant) | 0.079 | 0.055 | | 1.435 | 0.164 |
| | | | | Concentration | 0.068 | 0.095 | 0.142 | 0.715 | 0.481 |
| | male | *P. bras* | 1 | (Constant) | 0.089 | 0.109 | | 0.820 | 0.420 |
| | | | | Concentration | 0.376 | 0.188 | 0.371 | 1.996 | 0.057 |
| | | *P. mann* | 1 | (Constant) | 0.167 | 0.138 | | 1.209 | 0.244 |
| | | | | Concentration | 0.283 | 0.238 | 0.285 | 1.190 | 0.251 |
| | | *P. napi* | 1 | (Constant) | 0.311 | 0.119 | | 2.608 | 0.014 |
| | | | | Concentration | 0.366 | 0.206 | 0.318 | 1.776 | 0.087 |
| | | *P. rapa* | 1 | (Constant) | 0.121 | 0.060 | | 1.995 | 0.054 |
| | | | | Concentration | 0.185 | 0.114 | 0.269 | 1.626 | 0.113 |

**Table S9)** *Continued*

| | | | | | B | Std. Error | Beta | | |
|---|---|---|---|---|---|---|---|---|---|
| li | female | *P. bras* | 1 | (Constant) | 0.109 | 0.071 | | 1.536 | 0.133 |
| | | | | Concentration | 0.456 | 0.122 | 0.524 | 3.743 | **0.001** |
| | | *P. mann* | 1 | (Constant) | 0.630 | 0.346 | | 1.819 | 0.082 |
| | | | | Concentration | 0.468 | 0.597 | 0.165 | 0.784 | 0.441 |
| | | *P. napi* | 1 | (Constant) | 0.299 | 0.119 | | 2.513 | 0.017 |
| | | | | Concentration | 0.496 | 0.205 | 0.383 | 2.420 | **0.021** |
| | | *P. rapa* | 1 | (Constant) | 0.240 | 0.058 | | 4.156 | 0.000 |
| | | | | Concentration | 0.153 | 0.100 | 0.293 | 1.533 | 0.138 |
| | male | P. bras | 1 | (Constant) | 0.186 | 0.118 | | 1.573 | 0.128 |
| | | | | Concentration | 0.466 | 0.204 | 0.415 | 2.282 | **0.031** |
| | | P. mann | 1 | (Constant) | 0.426 | 0.181 | | 2.351 | 0.032 |
| | | | | Concentration | 0.361 | 0.313 | 0.278 | 1.156 | 0.265 |
| | | P. napi | 1 | (Constant) | 0.465 | 0.154 | | 3.026 | 0.005 |
| | | | | Concentration | 0.410 | 0.265 | 0.281 | 1.548 | 0.133 |
| | | P. rapa | 1 | (Constant) | 0.253 | 0.086 | | 2.937 | 0.006 |
| | | | | Concentration | 0.298 | 0.158 | 0.316 | 1.885 | 0.068 |
| me | female | P. bras | 1 | (Constant) | 0.137 | 0.054 | | 2.564 | 0.015 |
| | | | | Concentration | 0.391 | 0.092 | 0.571 | 4.236 | **0.000** |
| | | P. mann | 1 | (Constant) | 0.777 | 0.395 | | 1.965 | 0.062 |
| | | | | Concentration | 0.641 | 0.681 | 0.197 | 0.940 | 0.357 |
| | | P. napi | 1 | (Constant) | 0.323 | 0.105 | | 3.066 | 0.004 |
| | | | | Concentration | 0.488 | 0.181 | 0.419 | 2.687 | **0.011** |
| | | P. rapa | 1 | (Constant) | 0.178 | 0.067 | | 2.674 | 0.013 |
| | | | | Concentration | 0.369 | 0.115 | 0.541 | 3.214 | **0.004** |
| | male | P. bras | 1 | (Constant) | 0.331 | 0.122 | | 2.715 | 0.012 |
| | | | | Concentration | 0.593 | 0.210 | 0.492 | 2.825 | **0.009** |
| | | P. mann | 1 | (Constant) | 0.346 | 0.203 | | 1.710 | 0.107 |
| | | | | Concentration | 0.454 | 0.349 | 0.309 | 1.301 | 0.212 |
| | | P. napi | 1 | (Constant) | 0.425 | 0.148 | | 2.872 | 0.008 |
| | | | | Concentration | 0.451 | 0.255 | 0.317 | 1.768 | **0.088** |
| | | P. rapa | 1 | (Constant) | 0.161 | 0.069 | | 2.323 | 0.026 |
| | | | | Concentration | 0.360 | 0.129 | 0.438 | 2.799 | **0.009** |

#### S10 Interspecific variation and genetic basis of olfactory responses to post-mating odors (Figure 3)

##### *S10.1 Genome editing and full panel EAG results*

CRISPR/Cas9 was employed to disrupt the gene coding for olfactory receptor OR45b by targeting a region in the first exon with a single guide RNA. The potential off-target sites were searched by using bioinformatic tools (Exonerate 2.0 and online website CHOPCHOP v3) (Zhu et al., 2014; Labun et al., 2019). No off-target binding sites were found in the coding regions. Around 40 % of the injected eggs successfully hatched, and the caterpillars survived to butterflies. Sequenc-ing results showed that in one mutant line two base pairs were inserted in the first exon (Figure S4A). Transmembrane domain predictions showed that OR45b has a seven-transmembrane domain while the genome-edited OR45b exhibits an aborted coding sequence led by an early stop codon (Figure S3B). A couple of mutant butterflies with the same mutation were collected to generate homozygous mutants, each generation of *OR45b* KO insects was checked by PCR and DNA sequencing to ensure the homozygous colony. The offspring of these homozygous mutants was used in our experiments.

A panel of 47 volatile chemicals were employed to assess the electrophysiological responses of both wildtype and mutant butterflies of both sexes. We found that *OR45b* KO of both males and females showed reduced EAG response to seven chemicals, includ-ing the plant volatiles 1,8-cineole, benzaldehyde, limonene, *β*-caryophyllene, methyl salicylate and 3-hydroxy-2-butanone, as well as the anti-aphrodisiac pheromone benzyl cyanide (Figure S4C & D). In addition, only males have reduced antennal responses to another seven chemicals: (*E*)-2-hexenal, 2-methylbutanal, *α*-pinene, indole, benzo-thiazole and butyl isothiocyanate (ITC) (Figure S4C) and only females exhibited reduced responses to five chemicals (1-octen-3-ol, 3-octanol, phenylacetaldehyde, (*Z*)-2-pentenal, phenyl ITC) (Figure S4D).

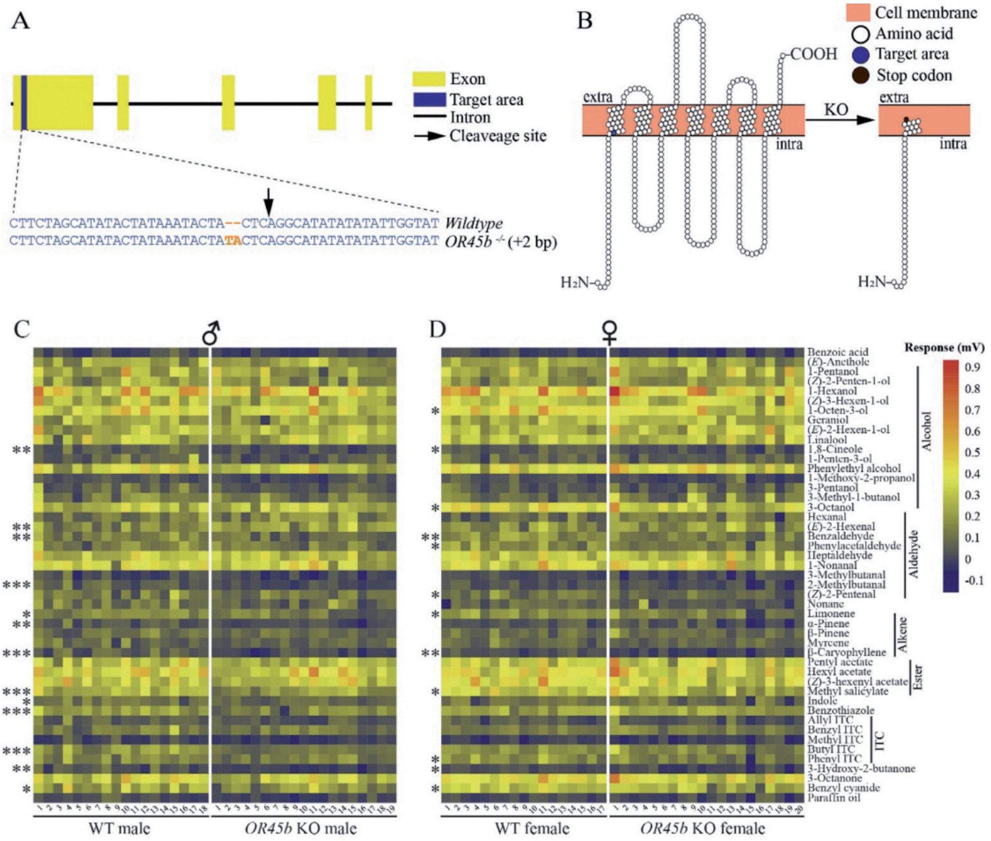


**Figure S4)** Odorant receptor gene *OR45b* editing and electroantennogram (EAG) responses in *Pieris brassicae* butterflies. (A) *OR45b* gene structure and mutation site. Yellow blocks indicate exons of *OR45b*, blue block indicates the area that was magnified and mutant sites were highlighted in bold orange. (B) *OR45b* gene transmembrane domain predictions in wildtype (WT) and knockout (KO) butterflies. Orange blocks indicate neuronal membrane; white circles indicate amino acids; blue circle indicates the target site; brown circle indicates the early stop codon; extra and intra are extracellular and intracellular. (C) EAG response of male butterflies from both butterfly genotypes. (D) EAG response of female butterflies from both butterfly genotypes. Chemical names are indicated on the right panel and classified into chemical classes. Significantly different response to chemicals between the two genotypes were indicated by asterisks * (0.01 < P < 0.05), ** (0.001 < P < 0.01) and *** (P < 0.001). Differences between WT and KO butterflies were tested by using a Student's t-test when the data was normally distributed and had similar variance or tested by using the Kruskal-Wallis test when the criteria of the Student's t-test did not apply for the data. The intensity of butterfly antennal responses is indicated by colors from dark blue (lowest) to red (highest), which is shown in the legend. N=17-20 butterflies of each sex for each genotype.

**Table S9)** EAG response for 5 compounds of interest in knockout versus wildtype *Pieris brassicae* (N= 17 – 20 butterflies)

| Odor | sex | test | p-value |
|---|---|---|---|
| Benzyl cyanide | Male | T-test | 0.0491 |
| | Female | T-test | 0.0242 |
| Methyl salycilate | Male | T-test | 0.0007 |
| | Female | T-test | 0.0461 |
| Indole | Male | T-test | 0.0149 |
| | Female | T-test | 0.6072 |
| Cis-3-hexynl acetate | Male | T-test | 0.05321 |
| | Female | Kruskal-wallis | 0.0511 |
| Linalool | Male | T-test | 0.7839 |
| | Female | T-test | 0.4082 |

#### S11 Functional consequences of post-mating odor perception and emission in Pieris brassicae. (Figure 4)

##### *S11.1 Butterfly egg-laying behavior and mating frequency*

To evaluate the effect of impaired olfaction on oviposition, the numbers of eggs laid by both butterfly genotypes were counted for seven days in experiment (1) where a couple of newly enclosed butterflies were placed in a cage and were allowed to mate freely. None of the butterflies deposited eggs on the first two days, Day 0 and Day 1. On Day 2 both genotypes of butterflies started laying eggs although only a few eggs were found on the plants and there was no difference between the two butterfly genotypes. From Day 3 onwards, *OR45b* KO butterflies laid significantly more eggs than WT butterflies (P < 0.0001, GLM negative binominal) (Figure S6). We further examined the number of eggs hatching between the genotypes: we found that *OR45b* KO butterflies laid significantly more fertilized eggs developing into caterpillars than WT butterflies (P = 0.0002, Wilcoxon rank-sum test), while egg-hatching rates were comparable between the two genotypes (P = 0.0620, Wilcoxon rank-sum test). Female butterflies were collected and dissected after the experiment. After dissection of the bursa copulatrix we found that *OR45b* KO butterflies tended to have mated significantly more times than WT butterflies (P = 0.0297, Wilcoxon rank-sum test). Interestingly, we found two full spermatophores in some of the *OR45b* KO female butterflies, indicating that mating occurred twice in quick succession, before the first spermatophore was fully or even partially digested (Figure S6).

We then compared the numbers of eggs laid by butterflies that had mated only once and did so for the comparison between the two butterfly genotypes in experiment (Figure S6). *OR45b* KO butterflies laid significantly more eggs on *B. oleracea* plants than WT butterflies since Day 1 (P < 0.0001, GLM negative binominal) (Figure 6.4B). The number of hatched eggs per female was similar for once-mated WT butterflies and once-mated *OR45b* KO butterflies (P = 0.5323, Student's t-test). However, the proportion of eggs that hatched was lower for *OR45b* KO butterflies than for WT butterflies (P = 0.0019, Wilcoxon rank-sum test).

In order to determine which sex determines the higher number of eggs laid by butterflies, wildtype and mutant butterflies were crossed and butterflies were allowed to mate freely, a cabbage plant was provided as egg-laying substrate in experiment 3. The *OR45b* KO females that mated with WT males laid significantly more eggs on cabbage than WT females that mated with KO males (P = 0.0015, GLM negative binomial) (Figure S6). WT females mated with KO males and KO females mated with WT males yielded a similar number of hatching eggs (P = 0.4238, Wilcoxon rank-sum test) (Figure S6) with a similar egg-hatching rate (P = 0.7318, Wilcoxon rank-sum test). Female butterflies of both genotypes were then collected and dissected; dissection results showed that the female butterflies in the two combinations had mated a similar number of times (P = 0.1360, Wilcoxon rank-sum test).

***S11.2 Butterfly host-plant selection and egg parasitoid preferences***

We investigated the behavioral responses of mated female butterflies to previously egg-infested and non-infested plants. WT female butterflies spent significantly more time before the first contact to the plants than *OR45b* KO butterflies (P = 0.0001, Wilcoxon rank-sum test) (Figure 6.4A). WT females were slightly more attracted to non-infested plants over egg-infested plants, although this difference was not significant (P = 0.1599), while *OR45b* KO females did not show any bias between the two plants (P = 0.4349); there is no significant difference in the final host-plant selection between the two genotypes (P = 0.1217, Chi-square test) (Figure S7).

Next, the behavioral preference of *T. evanescens* wasps was tested to evaluate the role of *OR45b* in tritrophic interactions (Figure 6.4C). The wasps significantly preferred mated female butterflies that were treated differently (P <0.0001, one-way ANOVA). In which, the wasps spent more time in the presence of *OR45b* KO mated females over the other butterflies (P = 0.0001, mated KO vs virgin KO; P = 0.0052, mated KO vs mated WT; P < 0.0001, mated KO vs virgin WT), while they did not prefer mated female WT butterflies over any of the genotypes of virgin butterflies (P = 0.2445, one-way ANOVA). To clarify why odors of mated *OR45b* KO female butterflies were more attractive to the egg parasitoids, we analyzed matings of both butterfly genotypes by dissecting the spermatophore. The mated female butterflies of both genotypes harbored a similar number of spermatophores (P = 0.1396, Wilcoxon rank-sum test) (Figure S9).

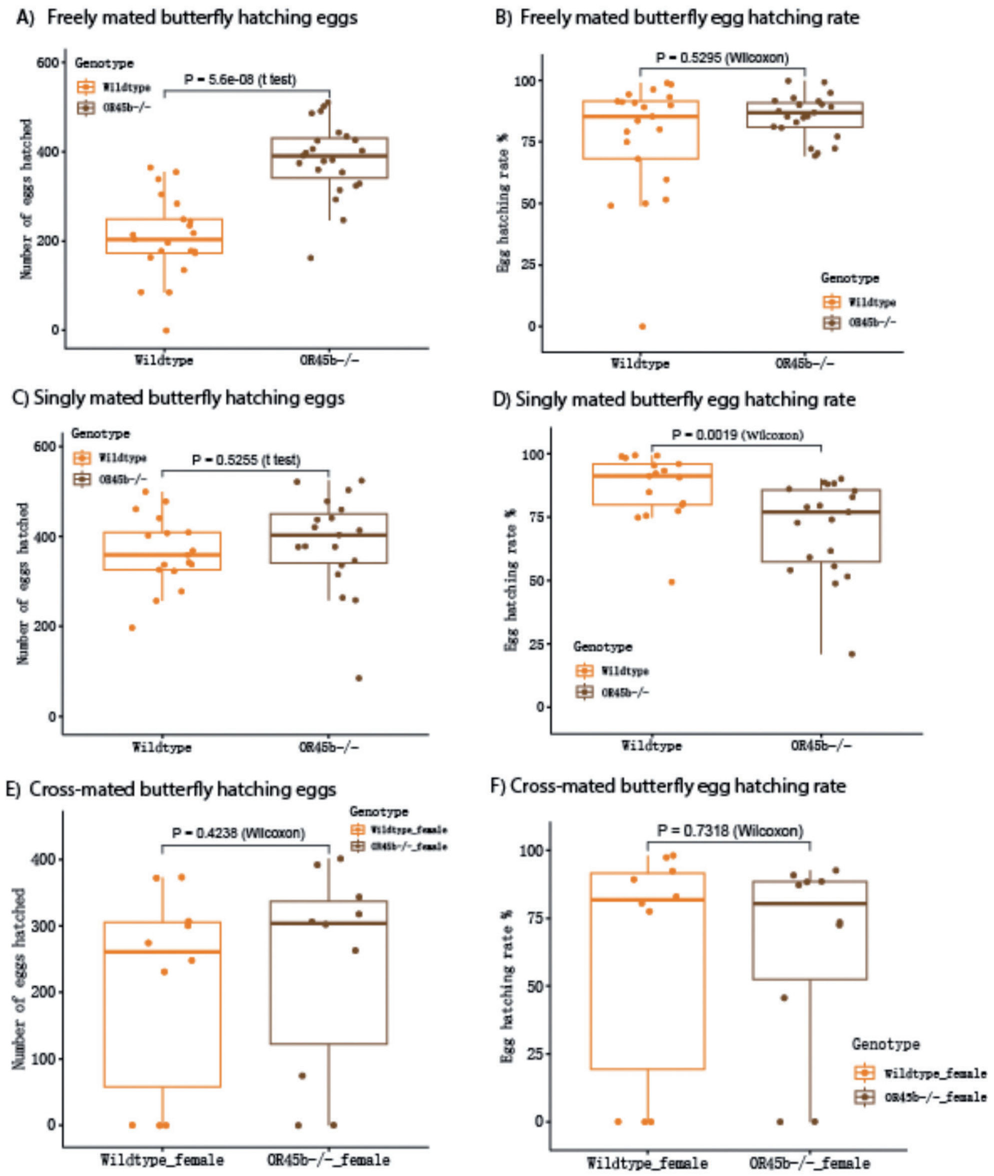


**Figure S6)** Comparison between wildtype (WT) and *OR45b* knockout (KO) *Pieris brassicae* in egg-laying behavior, number of matings and egg hatching success and rate. (A) Egg-laying dynamics of both genotypes of butterflies, a significant difference was detected with GLM with negative binomial distribution (n=14 for both genotypes). (B) Number of hatched eggs laid by both genotypes. Wilcoxon rank-sum test (n=14 for both genotypes). C) Number of eggs laid by females which mated one time. D) Number of eggs hatching, laid by females which mated one time. Egg-hatching rates by both genotypes (n=14 for both genotypes). E) Eggs laid in WT versus pair with only the female butterfly as KO but the male is WT. F) Hatching rate of eggs laid by WT versus pair where only the female is KO.

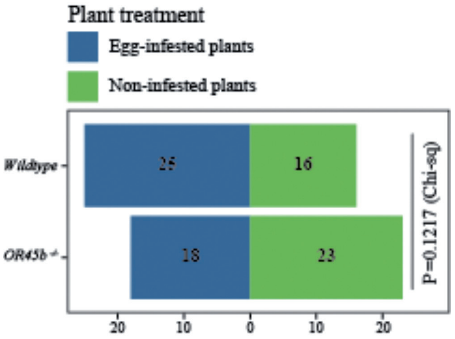


**Figure S7)** Oviposition choice of mutant and wildtype *Pieris brassicae*. Bars show the plants that both genotypes of female butterflies oviposited on. Blue and green indicate non-infested and egg-infested plants respectively. Difference was tested by Chi-square test.

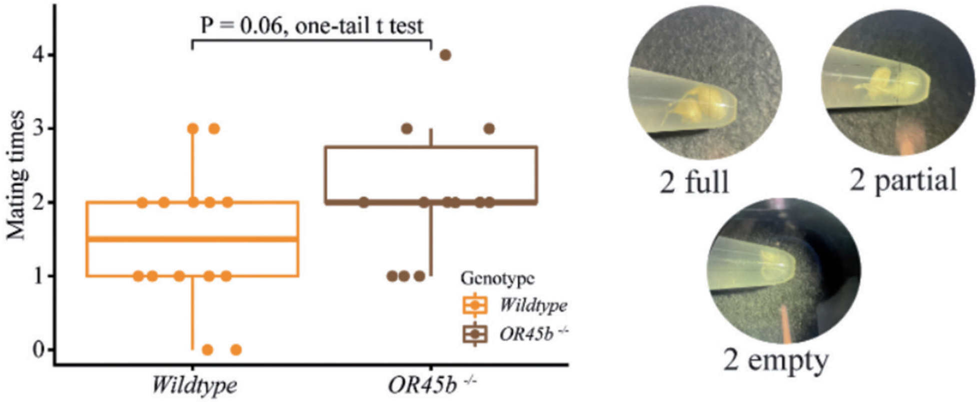


**Figure S8)** When allowed to mate freely over 10 days, knockout *Pieris brassicae* butterflies mated more often than wildtype butterflies and sometimes contained more than one full spermatophore at once.

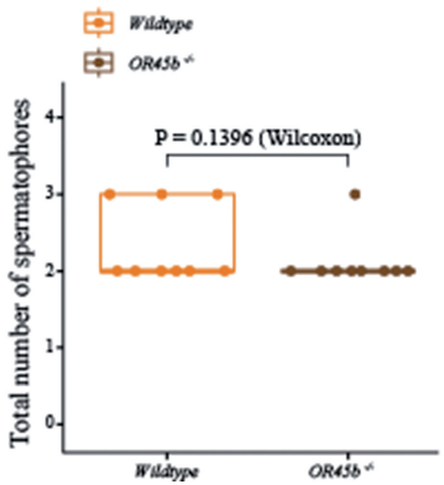


**Figure S9)** Similar mating rate of WT and KO *Pieris brassicae* used in the experiments where *Trichogramma* preference was tested in a static olfactometer.

**Table S10)** First choice of *Parus major* for virgin *Pieris brassicae* female butterflies, which were control (10 ul hexane treated) or treatment (10 ul hexane and benzyl cyanide). Results of binomial test are shown.

| | | Category | N | Observed Prop. | Test Prop. | Exact Sig. (2-tailed) |
|---|---|---|---|---|---|---|
| Choice_Trial1 | Group 1 | Control | 9 | .43 | .50 | .664 |
| | Group 2 | Treatment | 12 | .57 | | |
| | Total | | 21 | 1.00 | | |
| Choice_Trial2 | Group 1 | Control | 19 | .90 | .50 | <.001 |
| | Group 2 | Treatment | 2 | .10 | | |
| | Total | | 21 | 1.00 | | |
